# Trim28 plays an indispensable role in maintaining functions and transcriptional integrity of hematopoietic stem cells

**DOI:** 10.1101/2022.07.12.499765

**Authors:** Giovannino Silvestri, Chozha Vendan Rathinam

## Abstract

Hematopoiesis is a complex process that is regulated at multiple levels and through both intrinsic and extrinsic regulators. Trim28 is a multidomain protein that functions as a transcription factor and E3 ubiquitin ligase. Recent studies have established the pivotal roles of Trim28 in a variety of pathophysiologic processes including inflammation, autoimmune disorders, viral pathogenesis, cancer, tumor microenvironment and epithelial-to- mesenchymal transition. However, importance of Trim28 in early hematopoiesis, particularly in the maintenance and functions of hematopoietic stem and progenitor cells (HSPCs), remains largely unknown. In the present study, we identified Trim28 as a critical regulator of self-renewal, quiescence, and functions of hematopoietic stem cells (HSC). Conditional ablation of Trim28 in HSCs leads to perinatal lethality, due to pancytopenia within the hematopoietic lineage. Loss of Trim28 in HSCs affects their differentiation into myeloid-, erythroid-, and lymphoid- lineages. Trim28 deletion leads to altered frequencies and absolute numbers of HSCs, Multipotent Progenitors (MPPs) and lineage committed progenitors in the bone marrow (BM). Primary- and secondary- BM transplantation studies identified an indispensable and a cell-intrinsic role for Trim28 in HSCs. Mechanistic studies identified hyperproliferation of HSPCs and deregulated expression of cell cycle regulators in HSPCs. Strikingly, gene expression studies demonstrated that Trim28 deficiency impairs the transcriptional landscape and transcription factors signatures of HSPCs. In essence, these studies highlight a previously unknown role for Trim28 in the maintenance, functions, and multi-lineage differentiation of HSPCs.

## Introduction

Hematopoiesis is a process through which all blood/immune cells are generated in the primary lymphoid organs, such as the bone marrow (BM) and thymus and released into the bloodstream to circulate throughout the body. In response to a specific stimulus, the most primitive long-term (LT-) hematopoietic stem cells (HSCs) enter a proliferative phase and undergo either self-renewal or multi-lineage differentiation program^1,2^. Once homeostatic balance between myeloid and lymphoid differentiation is established in the periphery, HSCs return to an inactive quiescent or dormant phase^1,2^. Research over the past few decades, including our own, has identified a variety of cell-extrinsic and -intrinsic mechanisms in the regulation of self-renewal and differentiation of HSCs^3–6^. These include endogenous players, such as transcription factors, cell cycle regulators, microRNAs, cytokine receptors and E3 ubiquitin ligases, and exogenous stimuli, such as chemokines, hematopoietic cytokines, hormones, and inflammatory mediators^3–6^. Fine-tuning the levels of both intrinsic and extrinsic regulators is essential to maintain normal hematopoiesis. Indeed, deregulated mechanisms and pathways that alter expression of these molecular regulators result in pathologic hematopoiesis, including anemia, bone marrow failure, immunodeficiencies and cancer.

Tripartite Motif (Trim) family of proteins (~ 80 in humans) are highly conserved regulators of fundamental cellular processes in both vertebrates and invertebrates^7,8^. Most Trim members have a RING-finger domain, one or two Zinc-finger domains and its associated coiled-coil region on their N terminus and a COS domain, FNIII repeat, PRY domain, SPRY domain, acid-rich region, FIL domain, PHD domain, bromodomain, MATH domain, ARF domain and transmembrane region on their C terminus^7^. Trim proteins play pivotal roles in transcriptional regulation, cell proliferation, apoptosis, autophagy, DNA repair and oncogenesis through their unique capacities to perform dual functions as E3 ubiquitin ligases and transcriptional repressors/activators^9–11^. Indeed, deregulated expression of Trim family of genes causes several human diseases and disorders, such as cancer, immunological diseases, and developmental disorders^7,12,13^.

Trim28 (also known as kruppel-associated box-associated protein (Kap)-1 and transcriptional intermediary factor (Tif)- 1β) is expressed in the nucleus of most cell types and identified as a scaffolding protein without intrinsic repressive or DNA-binding properties^14,15^. Trim28 is a multi-functional protein and has been shown to play important roles in ubiquitylation ^16^ DNA damage^17^, establishment of genetic imprinting^18^, regulating transcription through RNA polymerase II promoter-pausing and pause release^19^, and retrotransposon silencing^15^.

Genetic inactivation of *Trim28* results in an arrest of embryonic development between day (E) 5.5 and 8.8^20^. Deletion of maternal *Trim28* causes embryonic lethality due to misregulation of genomic imprinting^21,22^ and male-predominant early embryonic lethality^23^. Systemic deletion of Trim28 in adult mice results in acute mortality^24^. Induction of Trim28 haploinsufficiency triggers bi-stable epigenetic obesity^25^ and affects selfrenewal of spermatogonial stem cells^24^. Trim28 has been shown to be indispensable for regulating pluripotency^26^ and self-renewal^27^ of embryonic stem cells, the formation of stable induced-pluripotent stem cells^28^ and somatic cell reprograming^29^. Within the hematopoietic system, functions of Trim28 are studied in B cells^30^, T cells^31–33^ and erythrocytes^34,35^. Of note, to date, significance of Trim28 in the generation, maintenance and functions of HSCs and hematopoietic progenitors remain largely unknown. To this end, in the present report, we studied the physiological roles of Trim28 in hematopoietic stem and progenitor cells (HSPCs). Our data indicated that conditional deletion of Trim28 in HSCs results in severe hematopoietic abnormalities, including multi-lineage hematopoietic defects, pancytopenia, and perinatal lethality.

## Results

### Trim28 prevents pancytopenia and perinatal lethality

To identify the functions of Trim 28 in HSCs, we conditionally ablated Trim28 in HSCs by crossing *Trim28*^F/F^ mice with Vav^cre/+^ transgenic mice to generate *Trim28*^F/F^ Vav^cre/+^ (henceforth referred to as “KO”) mice. Our previous studies established that crossing with Vav^cre^ deleter strain faithfully deletes floxed alleles in all (>99%) hematopoietic cells (including in LT-HSCs) of BM, spleen, and thymus, but not in non-hematopoietic cells/organs^36,37^. Indeed, our analysis indicated loss of Trim28 in the cells of BM and spleen (**Supplemental Figure 1A**). To identify if the KO mice are born at a mendelian ratio, we bred *Trim28*^F/F^ (female or male) mice with Trim28^F/+^ Vav^Cre/+^ (male or female) mice, respectively, and generated a total of 240 pups. Our genotyping studies identified ~ 30.42% of pups with Trim28^F/+^, ~ 28.33% of pups with Trim28^F/F^ and ~ 35.83% of pups with Trim28^F/+^ Vav^Cre/+^ genotypes (**Supplemental Figure 1 B-D**). However, only ~ 5.42% of pups were born with Trim28^F/F^ Vav^Cre/+^ genotype (**Supplemental Figure 1 B-D**), suggesting that the KOs were not born at mendelian ratio. Next, we studied the survival pattern of KO mice and our analysis indicated that ~ 83.3% of KO pups died between postnatal day (P) 5 and P20 (**Figure 1A**). Even though only ~ 16.6% of KO mice survived after P20, they all died by P45 (**Figure 1A**). Therefore, data for the present studies were obtained from KO survivors (and age/gender matched Trim28^F/+^ Vav^Cre/+^ control (henceforth referred to as “Con.”) mice) between 3 and 4 weeks after birth. Analysis of gross morphology of KO mice indicated a reduction of body size and weight (**Figure 1B and data not shown**). Complete blood count (CBC) analysis indicated reduced numbers of total WBCs, lymphocytes, neutrophils, RBCs and platelets in the peripheral blood of KO mice (**Figure 1C**). Together, these data suggested that loss of Trim28 results in perinatal lethality, possibly due to multi-lineage hematopoietic defects and pancytopenia.

**Figure 1.**
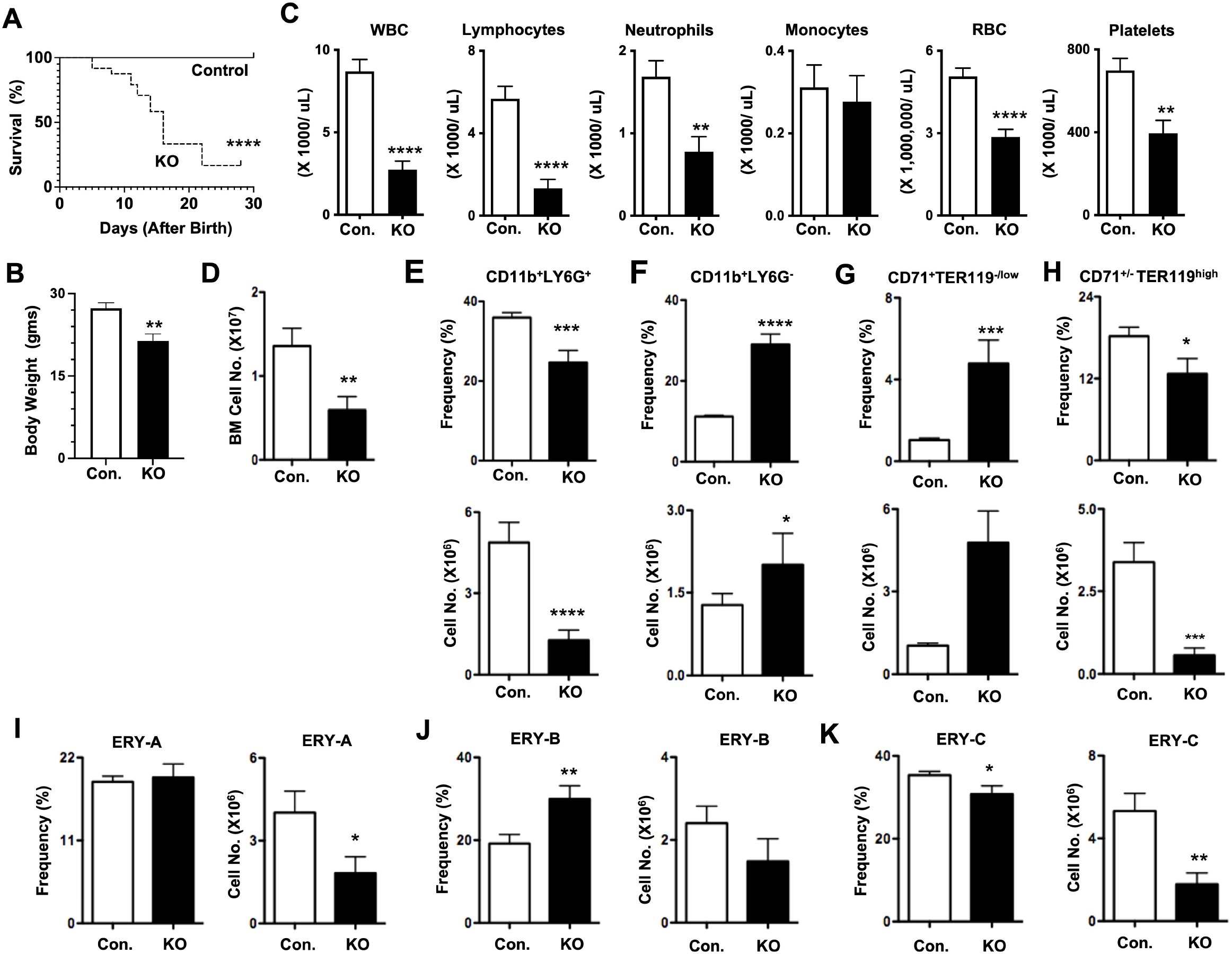
Trim28 deficiency leads to premature death & defective myeloid differentiation. **A.** Kaplan-Meier survival curve analysis of Trim28^F/F^ Vav^Cre^ (KO) and Control (Con.) mice (n=24). Significance (****, P < 0.0001) was assessed using the log-rank test. **B.** Body weight of KO and Con. mice after 3 weeks of birth (n= 5). **C.** Complete Blood Count (CBC) analysis of KO (n=19) and Con. (n=20) mice between 3 and 4 weeks of birth. **D.** Total cell counts of bone marrow (BM) of KO (n=15) and Con. (n=20) mice between 3 and 4 weeks of birth. **E-H.** Frequencies (**top**) and absolute numbers (**bottom**) of granulocytes (**E**), myelo/monocytes (**F**), proerythroblasts (**G**) and total erythroblasts (**H**) in the BM of KO (n=21) and Con. (n=26) mice between 3 and 4 weeks of birth. **I-K.** Frequencies (**left**) and absolute numbers (**right**) of Immature erythroblasts (EryA) (**I**), less-mature erythroblasts (EryB) (**J**), and most-mature erythroblasts (EryC) (**K**) in the BM of KO (n=14) and Con. (n=19) mice between 3 and 4 weeks of birth. All data represent mean ± SEM. Two-tailed Student’s *t* tests were used to assess statistical significance (*, P < 0.05; **, P < 0.01; ***, P < 0.001; ****, P < 0.0001).

### Trim28 regulates differentiation of myeloid, erythroid and lymphoid cells

To study the functions of Trim28 in the early differentiation pathways of the hematopoietic lineages, we analyzed the BM, spleen, and thymus of KO mice. Our cell count analysis indicated reduced absolute numbers of BM of KO mice (**Figure 1D**). Immunophenotyping studies revealed a reduction in both frequencies and absolute numbers of Gr1^+^CD11b^+^ granulocytes (**Figure 1E**) and an increase in frequencies and absolute numbers of Gr1^-^CD11b^+^ myelo/monocytic cells (**Figure 1F**) in the BM of KO mice. Analysis of erythroid lineage^38^ in the BM of KO mice indicated increased frequencies and normal absolute numbers of CD71^+^Ter119^low^ proerythroblasts (ProE) (**Figure 1G**) and reduced frequencies and absolute numbers of CD71^high/-^Ter119^high^ total eyrthroblasts (**Figure 1H**). Further analysis of erythroblasts^38^ indicated normal frequencies and absolute numbers of CD71^high^Ter119^high^FSc^high^ immature erythroblasts (EryA fraction) (**Figure 1I**), increased frequencies, but normal absolute numbers, of CD71^high^Ter119^high^FSc^low^ less-mature erythroblasts (EryB fraction) (**Figure 1J**) and reduced frequencies and absolute numbers of CD71^low^Ter119^high^FSc^low^ mature erythroblasts (EryC fraction) (**Figure 1K**) in the BM of KO mice. Finally, we studied B cell differentiation in the BM of KO mice. Our studies indicated normal frequencies, but reduced absolute numbers, of CD19^-^B220^+^ B-lineage progenitors (**Figure 2A**) and remarkably reduced frequencies and absolute numbers of CD19^+^B220^+^ B cells (**Figure 2B**) in the BM of KO mice. Overall, these data suggested that the differentiation of myeloid-, erythroid- and B-lineages in the BM is perturbed in the absence of Trim28.

**Figure 2.**
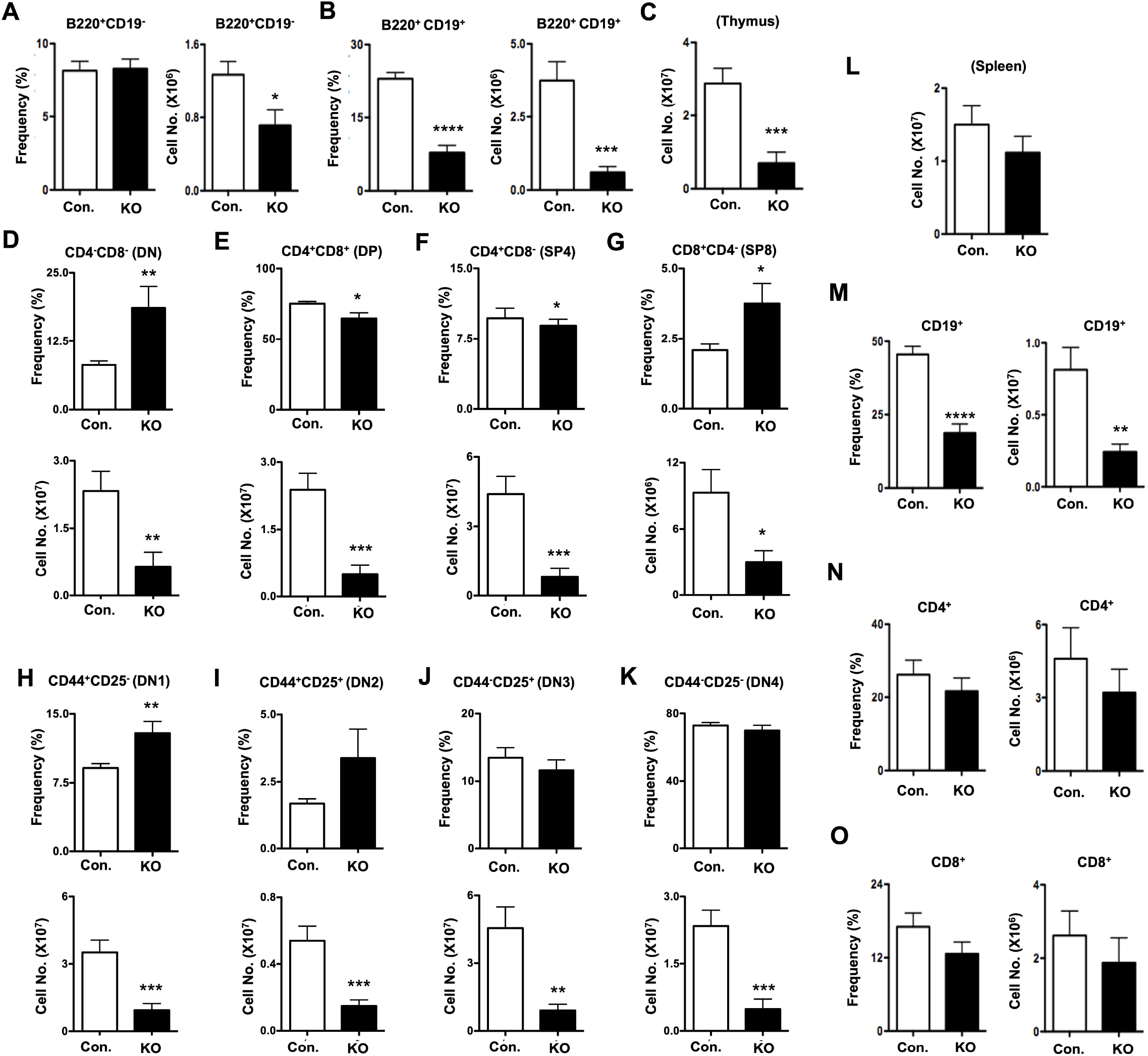
Loss of Trim28 results in diminished lymphoid differentiation. **A-B.** Frequencies (**left**) and absolute numbers (**right**) of pro-B cells (**A**) and pre-/immature-/ mature-B cells (**B**) in the BM of KO (n=16) and Con. (n=21) mice between 3 and 4 weeks of birth. **C.** Total cell counts of thymus from KO (n=16) and Con. mice (n=19) between 3 and 4 weeks of birth. **D-G.** Frequencies (**top**) and absolute numbers (**bottom**) of DN (**D**), DP(**E**), SP4 (**F**) and SP8 (**G**) subsets of thymus of KO (n=25) and Con. mice (n=30) between 3 and 4 weeks of birth. **H-K.** Frequencies (**top**) and absolute numbers (**bottom**) of DN1 (**H**), DN2(**I**), DN3 (**J**) and DN4 (**K**) fractions within DN subset of thymus from KO (n=25) and Con. mice (n=30) between 3 and 4 weeks of birth. **L.** Total cell counts of spleen from KO (n=15) and Con. mice (n=20) between 3 and 4 weeks of birth. **M-O.** Frequencies (**left**) and absolute numbers (**right**) of B cells (**M**), CD4^+^ T cells (**N**) and CD8^+^ T cells (**O**) in the spleen of KO (n=21) and Con. mice (n=26) between 3 and 4 weeks of birth. All data represent mean ± SEM. Two-tailed Student’s *t* tests were used to assess statistical significance (*, P < 0.05; **, P < 0.01; ***, P < 0.001; ****, P < 0.0001).

To assess the role of Trim28 in the development of T cells, we analyzed the thymus of KO mice. Our data indicated a remarkable reduction in the total counts of thymocytes in KO mice (**Figure 2C**). Immunophenotyping analysis of specific developmental stages of T cells in the thymus of KO mice indicated; an increase in frequencies, but reduced absolute numbers, of CD4-CD8- (double negative (DN)) thymocytes (**Figure 2D**); a decrease in frequencies and absolute numbers of CD4^+^CD8^+^ (double positive (DP)) thymocytes (**Figure 2E**); a decrease in frequencies and absolute numbers of CD4^+^CD8^-^ (single positive (SP) 4) thymocytes (**Figure 2F**); and increased frequencies, but reduced absolute numbers, of CD4^-^CD8^+^ (single positive (SP) 8) thymocytes (**Figure 2G**). Further division of DN subsets into specific developmental (DN1-4) stages suggested increased frequencies and reduced absolute numbers of CD44^+^CD25^-^DN1 subset (**Figure 2H**), and normal frequencies & reduced absolute numbers of CD44^+^CD25^+^ DN2 (**Figure 2I**), CD44^-^CD25^+^ DN3 (**Figure 2J**) and CD44^-^CD25^-^DN4 (**Figure 2K**) subsets in the thymus of KO mice. These data demonstrate that Trim28 functions are critical for the differentiation of T cells in the thymus.

Finally, we assessed the frequencies of myeloid, erythroid and lymphoid cells in the spleen of KO mice. Our data indicated normal splenic cell counts (**Figure 2L**), remarkably reduced frequencies and absolute numbers of B cells (**Figure 2M**), normal frequencies and absolute numbers of CD4^+^ (**Figure 2N**) and CD8^+^ (**Figure 2O**) T cells in the spleen of KO mice. Analysis of myeloid lineage indicated reduced frequencies, but normal, absolute numbers of Gr1^+^CD11b^+^ granulocytes (**Supplemental Figure 2A**), and increased frequencies, but normal absolute numbers, of Gr1^-^CD11b^+^ myelo/monocytic cells (**Supplemental Figure 2B**) in the spleen of KO mice. Erythroid lineage analysis of spleen from KO mice indicated; normal frequencies and absolute numbers of ProE (**Supplemental Figure 2C**); increased frequencies and absolute numbers of total erythroblasts (**Supplemental Figure 2D**); normal frequencies and absolute numbers of EryA (**Supplemental Figure 2E**) and EryB (**Supplemental Figure 2F**) fractions; and normal frequencies, but reduced absolute numbers, of EryC (**Supplemental Figure 2G**) fraction. Together, these data specify that the maintenance of myeloid-, erythroid-, and T-lineage in the peripheral lymphoid organ is modestly affected by the loss of Trim28. However, maintenance of peripheral B cells is strictly dependent on Trim28.

### Trim28 acts as a gatekeeper of HSPC pool in the BM

To identify the impact of Trim28 deficiency in the maintenance of HSPCs, we analyzed the BM of KO mice. Immunophenotyping analysis of lineage (Lin)^-^ undifferentiated cells and Lin^-^ Sca1^+^c-Kit^+^ (LSK) HSPCs indicated normal frequencies and absolute numbers in the BM of KO mice (**Figure 3A-B**). Further discrimination of LSK cells into distinct HSPC subsets^39^ from the BM of KO mice revealed; increased frequencies, but normal absolute numbers, of CD150^+^CD48^-^Flt3^-^LSK (LT-HSCs) (**Figure 3C**); strikingly decreased frequencies and absolute numbers of CD150^-^CD48^-^Flt3^-^LSK short-term (ST)-HSCs (**Figure 3D**); increased frequencies, but normal absolute numbers, of CD150^+^CD48^+^Flt3^-^LSK multi-potent progenitor (MPP) 2 (**Figure 3E**); and reduced frequencies and absolute numbers of CD150^-^CD48^+^Flt3^-^LSK MPP3 (**Figure 3F**) & Flt3^+^LSK MPP4 (**Figure 3G**) subsets.

**Figure 3.**
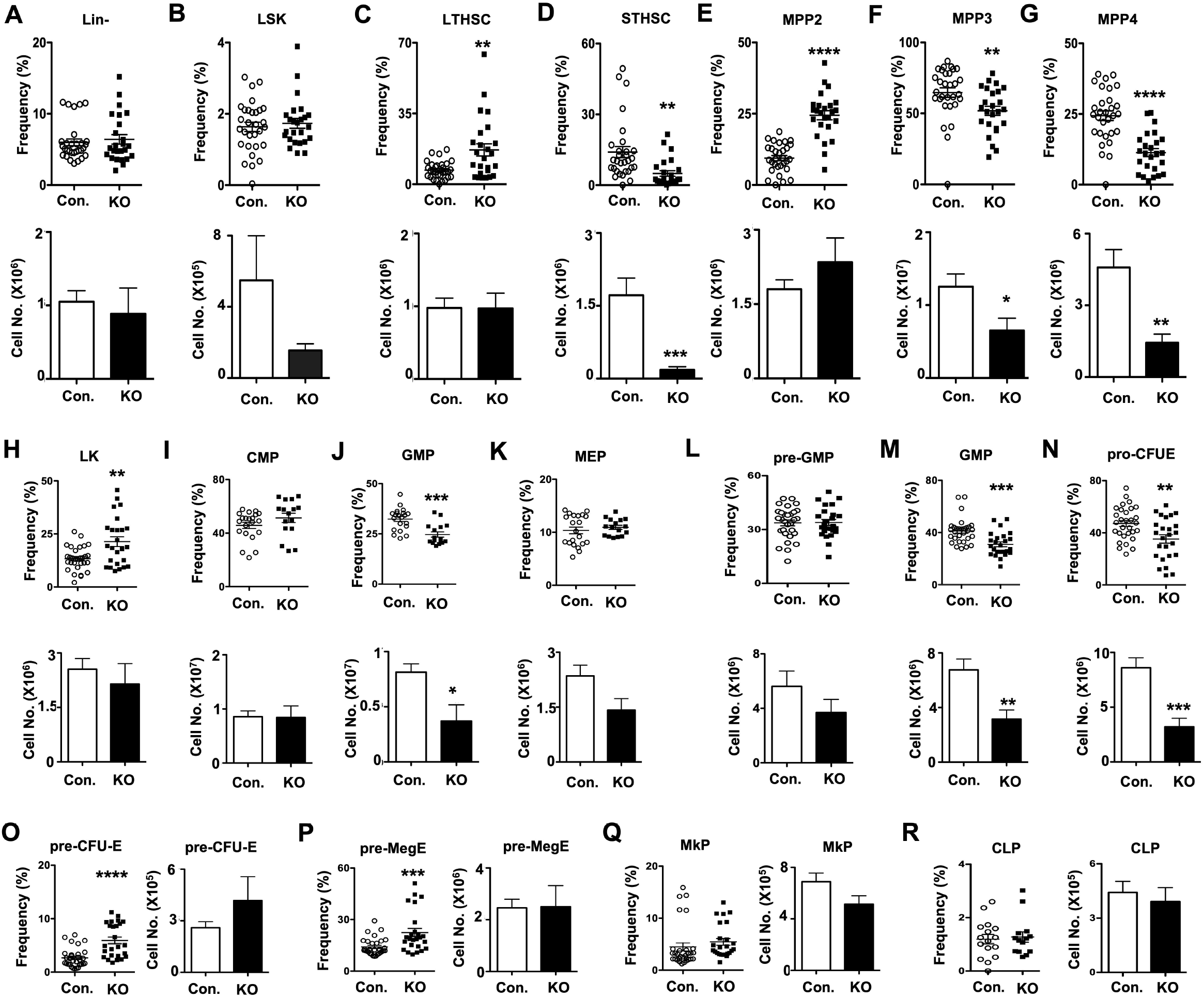
Lack of Trim28 alters HSC & progenitor pool of the BM. **A-B.** Frequencies (**top**) and absolute numbers (**bottom**) of Lin- (**A**) and LSK (**B**) cells in the BM of KO (n=25) and Con. mice (n=30) between 3 and 4 weeks of birth. **C-G.** Frequencies (**top**) and absolute numbers (**bottom**) of LT-HSC (**C**), ST-HSC (**D**), MPP2 (**E**), MPP3 (**F**) and MPP4 (**G**) subsets within LSK fraction of the BM from KO (n=25) and Con. mice (n=30) between 3 and 4 weeks of birth. **H.** Frequencies (**top**) and absolute numbers (**bottom**) of LK subset of the BM from KO (n=25) and Con. mice (n=30) between 3 and 4 weeks of birth. **I-K.** Frequencies (**top**) and absolute numbers (**bottom**) of CMP KO (n=16) and Con. (n=21) mice (**I**), GMP KO (n=16) and Con. (n=21) Mice (**J**) and MEP KO (n=16) and Con. (n=21) mice (**K**) subsets within LK fraction of the BM from KO and Con. mice between 3 and 4 weeks of birth. **L-N.** Frequencies (**top**) and absolute numbers (**bottom**) of pre-GMP (**L**), GMP (**M**) and pro-CFU-E (**N**) subsets within LK fraction of the BM from KO (n=25) and Con. mice (n=30) between 3 and 4 weeks of birth. **O-Q.** Frequencies (**left**) and absolute numbers (**right**) of pre-CFU-E (**O**), pre-MegE (**P**), MkP (**Q**) subsets within LK fraction of the BM from KO (n=25) and Con. mice (n=30) between 3 and 4 weeks of birth. **R.** Frequencies (**top**) and absolute numbers (**bottom**) of CLP within the BM of KO (n=14) and Con. mice (n=16) between 3 and 4 weeks of birth. All data represent mean ± SEM. Two-tailed Student’s *t* tests were used to assess statistical significance (*, P < 0.05; **, P < 0.01; ***, P < 0.001; ****, P < 0.0001).

To assess if the loss of Trim28 alters committed progenitors of specific hematopoietic lineage, we analyzed progenitors of the myeloid lineage^40^. Analysis of Lin^-^c-Kit^+^ (LK) total myeloid progenitors suggested an increase in frequencies, but normal absolute numbers, in the BM of KO mice (**Figure 3H**). Further division of LK subset indicated normal frequencies and relative numbers of CD34^+^CD16/32^-^LK common myeloid progenitors (CMPs) (**Figure 3I**) and CD34^-^CD16/32^-^LK megakaryocyte erythrocyte progenitors (MEPs) (**Figure 3K**) in the BM of KO mice. On the other hand, both frequencies and absolute numbers of CD34^+^CD16/32^+^LK granulocyte monocyte progenitors (GMPs) were reduced in the BM of KO mice (**Figure 3J**). To independently validate these findings, we followed an alternative and refined immunophenotyping strategy that distinguishes progenitors of the erythroid and megakaryocytic lineages^41^. Accordingly, our studies identified; reduced frequencies and absolute numbers of CD41^-^CD16/32^+^CD150^-^LK GMPs (**Figure 3L**); normal frequencies and absolute numbers of CD41^-^CD16/32^-^CD150^-^CD105^-^LK pre-GM (**Figure 3M**); reduced frequencies and absolute numbers of CD41^-^CD16/32^-^CD150^+^CD105^-^LK pro-erythrocytes + colony forming unit-erythrocytes (CFU-E) (**Figure 3N**); increased frequencies, but normal numbers of CD41^-^CD16/32^-^CD150^+^CD105^+^LK pre-CFU-E (**Figure 3O**) and CD41^-^CD16/32^-^CD150^+^CD105^-^LK pre-megakaryocyte and erythrocyte (pre-MegE) progenitors (**Figure 3P**); and normal frequencies and absolute numbers of CD41^+^CD150^+^LK megakaryocyte progenitors (MkP) (**Figure 3Q**) in the BM of KO mice. Finally, immunophenotyping studies confirmed normal frequencies and absolute numbers of IL7Ra^+^ Lin^-^ Sca1^+^c-Kit^low/-^ common lymphoid progenitors (CLPs) in the BM of KO mice (**Figure 3R**). Overall, these data highlight that Trim28 functions are essential for proper maintenance of HSC and MPP pool and their differentiation into myeloid- and erythroid-lineage committed progenitors.

### Trim28 has an indispensable and cell-intrinsic roles in HSC functions

To evaluate the role of Trim28 in regulating HSC functions, we performed bone marrow transplantation (BMT) studies. RBC depleted total BM cells of either control or KO mice were transplanted into lethally irradiated wildtype congenic recipients and donor derived hematopoiesis was studied. Analysis of peripheral blood after 4 weeks (**Figure 4A)** and 24 weeks (**Figure 4B)** of transplantation revealed reduced frequencies of CD45.2^+^ donor derived hematopoiesis and increased frequencies of CD45.1^+^ recipient derived hematopoiesis in recipients that received KO BM. Next, we assessed donor frequencies in the BM (**Figure 4C)** and spleen (**Figure 4D)** of recipients after 24 weeks of transplantation and data indicated reduction of CD45.2^+^ derived hematopoiesis in recipients that received KO BM.

**Figure 4.**
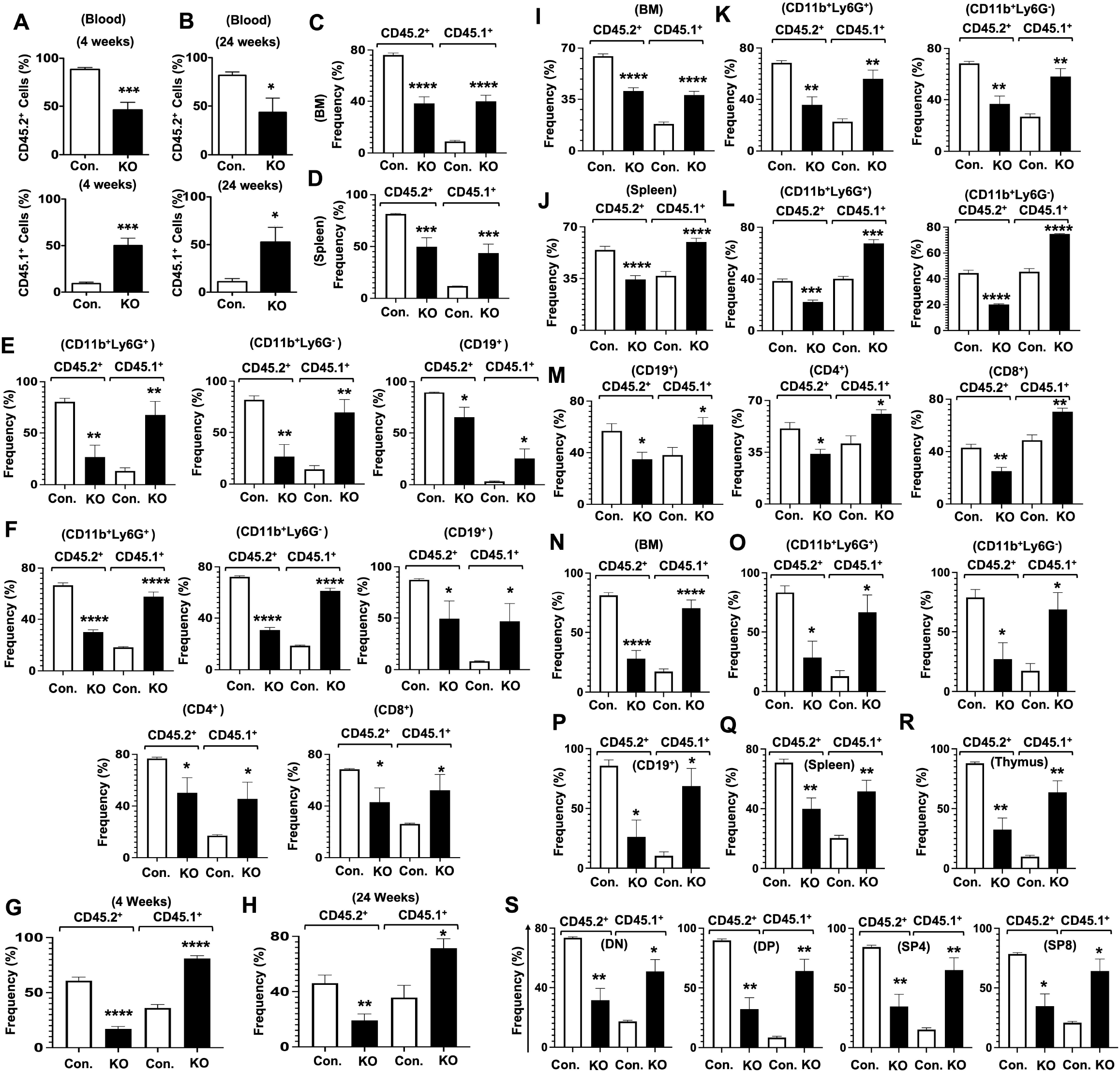
Deficiency of Trim28 suppresses HSC functions in a cell-intrinsic manner. **A-B.** Frequencies of donor- (CD45.2^+^) (**top**) and recipient- (CD45.1^+^) (**bottom**) derived hematopoiesis in the peripheral blood of wildtype (WT) recipients, after 4 weeks (**A**) KO (n=5) and Con. mice (n=5) and 24 weeks KO (n=4) and Con. mice (n=5) (**B**) of transplantation. **C-D.** Frequencies of donor- (CD45.2^+^) and recipient- (CD45.1^+^) derived hematopoiesis in the BM (**C**) and spleen (**D**) of WT recipients (n= 9-10), after 24 weeks of transplantation. **E.** Frequencies of donor- (CD45.2^+^) and recipient- (CD45.1^+^) derived granulocytes (**left**), myelo/monocytes (**middle**) and B cells (**right**) in the BM of WT recipients (n= 9), after 24 weeks of transplantation. **F.** Frequencies of donor- (CD45.2^+^) and recipient- (CD45.1^+^) derived granulocytes (**top-left**), myelo/monocytes (**top-middle**), B cells (**top-right**), CD4^+^ T cells (**bottom-left**) and CD8^+^ T cells (**bottom-right**) in the spleen of WT recipients (n= 9), after 24 weeks of transplantation. **G-H.** Frequencies of donor- (CD45.2^+^) and recipient- (CD45.1^+^) derived hematopoiesis in the peripheral blood of WT recipients (n= 14), after 4 weeks (**G**) and 24 weeks (**H**) of mixed-BMT. **I-J.** Frequencies of donor- (CD45.2^+^) and recipient- (CD45.1^+^) derived hematopoiesis in the BM (**I**) and spleen (**J**) of WT recipients (n= 14), after 24 weeks of transplantation. **K-L.** Frequencies of donor- (CD45.2^+^) and recipient- (CD45.1^+^) derived granulocytes (**left**) **and** myelo/monocytes (**right**) in the BM (**K**) and spleen (**L**) of WT recipients (n= 14), after 24 weeks of mixed-BMT. **M.** Frequencies of donor- (CD45.2^+^) and recipient- (CD45.1^+^) derived B cells (**left**), CD4^+^ T cells (**middle**), CD8^+^ T cells (**right**) in the spleen of WT recipients (n= 14), after 24 weeks of mixed-BMT. **N.** Frequencies of donor- (CD45.2^+^) and recipient- (CD45.1^+^) derived hematopoiesis in the BM of WT secondary-BMT recipients (n= 6), after 16 weeks of transplantation. **O-P.** Frequencies of donor- (CD45.2^+^) and recipient- (CD45.1^+^) derived granulocytes (**O; left**), myelo/monocytes (**O; right**), and B cells (**P**) in the BM of WT recipients (n= 6), after 16 weeks of secondary-BMT. **Q-R.** Frequencies of donor- (CD45.2^+^) and recipient- (CD45.1^+^) derived hematopoiesis in the spleen (**Q**) and thymus (**R**) of WT secondary-BMT recipients (n= 6), after 16 weeks of transplantation. **S.** Frequencies of donor- (CD45.2^+^) and recipient- (CD45.1^+^) derived DN (**far left**), DP (**left**), SP4 (**right**) and SP8 (**far right**) subsets of thymus in WT secondary-BMT recipients (n= 6), after 16 weeks of transplantation. **E, F, K, L, M, O, P & S**. Immune subsets were pre-gated based on their immunophenotypes and the frequencies of CD45.2^+^ vs. CD45.1^+^ cells within pre-gated fractions were calculated. All data represent mean ± SEM. Two-tailed Student’s *t* tests were used to assess statistical significance (*, P < 0.05; **, P < 0.01; ***, P < 0.001; ****, P < 0.0001).

Multi-lineage analysis in primary recipients that received KO BM suggested normal frequencies of granulocytes (Ly6G^+^CD11b^+^), myelo/monocytes (Ly6G^-^CD11b^+^) and B cells (CD19^+^) cells in the BM (**Supplemental Figure 3A**), and of granulocytes, myelo/monocytes, B cells, CD4^+^ T cells and CD8^+^ T cells within the spleen (**Supplemental Figure 3B**). However, analysis of CD45.2 vs. CD45.1 derived chimera within individual myeloid and lymphoid subsets indicated reduced CD45.2^+^ (KO) derived, whereas increased CD45.1^+^ (wildtype) derived, granulocytes, myelo/monocytes, and B cells within the BM of recipients that received KO BM (**Figure 4E**). Consistently, frequencies of CD45.2^+^ KO derived granulocytes, myelo/monocytes, B cells, CD4 T cells and CD8 T cells were diminished in the spleen of wildtype recipients (**Figure 4F**).

To test the capacities of Trim28 deficient HSCs to generate multi-lineage hematopoiesis under competitive settings and to assess if the compromised functions of Trim28 deficient HSCs are caused by cell intrinsic mechanisms, we performed mixed BMT studies. RBC depleted BM cells of either control or KO mice (CD45.2^+^) were mixed with BM cells of wildtype congenic mice (CD45.1^+^) at 1:1 ratio and transplanted into lethally irradiated wildtype congenic (CD45.1^+^) recipients. Analysis of peripheral blood after 4 weeks (**Figure 4G**) and 24 weeks (**Figure 4H**) of transplantation indicated a striking reduction of KO (CD45.2^+^) derived hematopoiesis within wildtype recipients. As expected, analysis of BM (**Figure 4I**) and spleen (**Figure 4J**) from mixed BMT recipients after 24 weeks of transplantation suggested reduced frequencies of CD45.2^+^ KO derived hematopoiesis. Multi-lineage analysis revealed normal frequencies of myeloid and lymphoid lineage cells within the BM (**Supplemental Figure 3C**) and spleen (**Supplemental Figure 3D**) of recipients that received KO + wildtype BM, when compared with Con + wildtype BM. Nevertheless, frequencies of KO (CD45.2^+^) derived granulocytes and myelo/monocytes within in BM (**Figure 4K**) and spleen (**Figure 4L**), and B cells, CD4^+^ T cells and CD8^+^ T cells within spleen (**Figure 4M**) were reduced in recipients that received KO + wildtype BM.

Finally, we performed secondary BMT studies using the BM of primary recipients (after 24 weeks of primary transplantation) from the total-BMT cohort. Analysis of recipients after 16 weeks of secondary transplantation exhibited a striking reduction of CD45.2^+^ KO derived overall hematopoiesis in the BM (**Figure 4N**). Multi-lineage analysis indicated diminished frequencies of CD45.2^+^ KO derived granulocytes, myelo/monocytes (**Figure 4O**) and B cells (**Figure 4P**) in the BM. Consistently, analysis of spleen (**Figure 4Q**) and thymus (**Figure 4R**) indicated reduced KO derived hematopoiesis in secondary recipients. Detailed analysis of thymus of secondary recipients suggested reduced frequencies of CD45.2^+^ KO derived DN, DP, SP4 and SP8 subsets (**Figure 4S**). In toto, data of BMT studies indicated that Trim28 deficiency leads to compromised HSC functions, in a cell intrinsic manner.

### Trim28 suppresses HSPC proliferation

To understand cellular mechanisms that cause diminished HSC functions in the absence of Trim28, we assessed the proliferation potential of HSPCs. Indeed, there exists a tight balance between quiescence and proliferation in HSCs and even minor perturbations on this key mode of HSC control can lead to premature exhaustion of HSC pool and functions^1–6^. To quantify proliferation of HSPCs subsets, we injected *i.p*. control and KO mice with Bromodeoxyuridine (BrdU) and analyzed after 24 hours, as we described earlier^42^. Our BrdU studies indicated an augmented proliferation (based on BrdU incorporation) of Lin- (**Figure 5A**), LSK (**Figure 5B**), LT-HSCs (**Figure 5C**), ST-HSCs (**Figure 5D**), MPPs (**Figure 5E**) and LK (**Figure 5F**) subsets of the KO BM.

**Figure 5.**
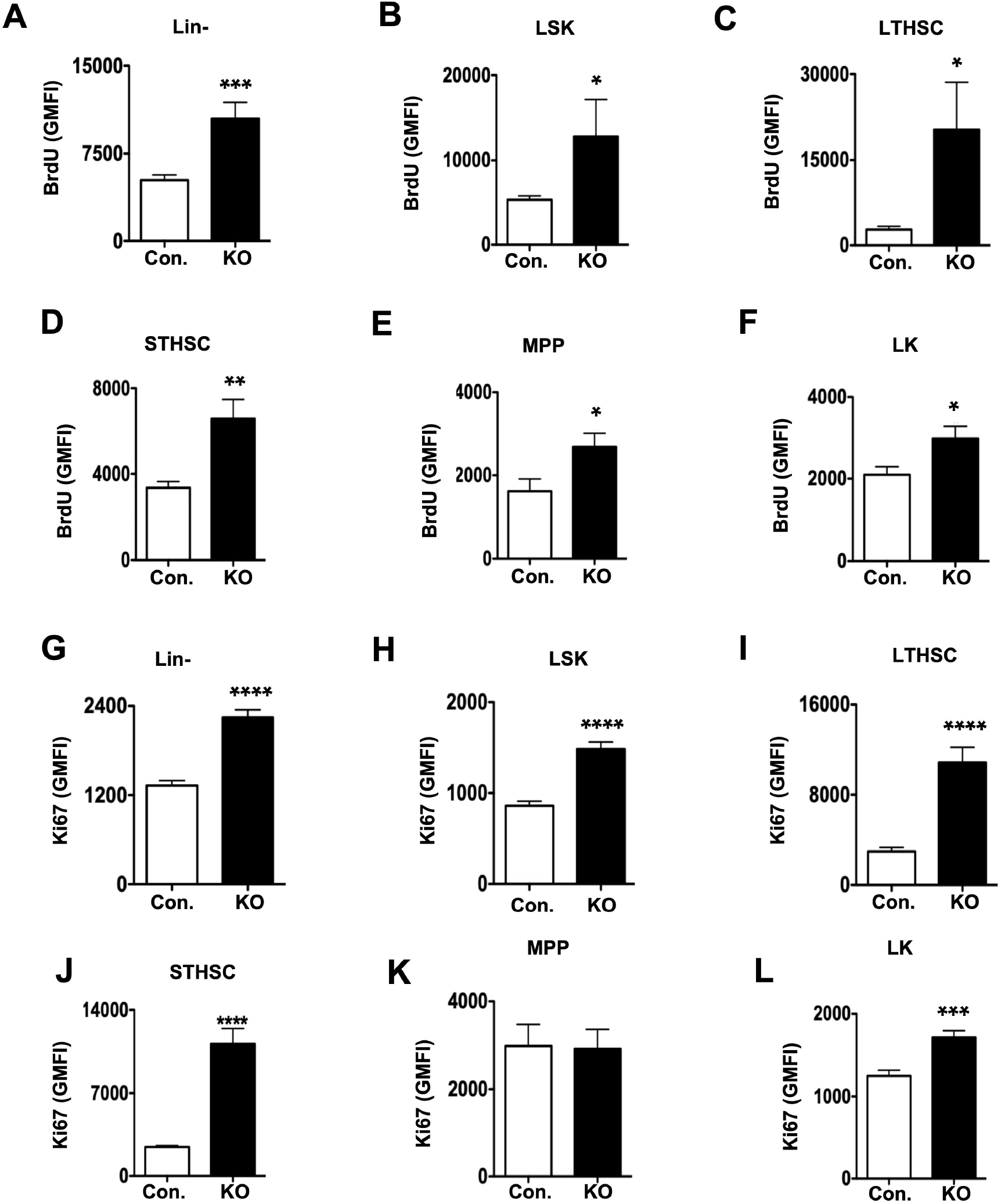
Ablation of Trim28 leads to hyperproliferation of HSPCs. **A-F.** Geomean fluorescence Intensity (GMFI) for BrdU^+^ of Lin- (**A)**, LSK (**B)**, LT-HSC (**C),** ST-HSC (**D), MPP** (**E)** and LK (**F) subsets** of the BM from KO (n=4-12) and control (n=6-16) mice. **G-L.** Geomean fluorescence Intensity (GMFI) for Ki67^+^ of Lin- (**G)**, LSK (**H)**, LT-HSC (**I),** ST-HSC (**J), MPP** (**K)** and LK (**L) subsets** of the BM from KO (n=4-12) and control (n=6-16) mice. Data shown in **A, B, C, F, G, H, K** & **L** are pool of two independent experiments, whereas data shown in **D, E, I** & **J** are representative of two independent experiments. All data represent mean ± SEM. Two-tailed Student’s *t* tests were used to assess statistical significance (*, P < 0.05; **, P < 0.01; ***, P < 0.001; ****, P < 0.0001).

To corroborate these findings through an alternative method, we measured nuclear Ki67 levels by flow cytometry^42^. Data of these studies specified that Ki67 levels were remarkably elevated in Lin- (**Figure 5G**), LSK (**Figure 5H**), LT-HSCs (**Figure 5I**), ST-HSCs (**Figure 5J**) and LK (**Figure 5L**) subsets of the KO BM.

To gain molecular insights into mechanisms that cause elevated proliferation of HSPCs in the absence of Trim28, we performed gene expression studies. We analyzed expression levels of positive regulators (Cyclins) and negative regulators (cyclin-dependent kinase inhibitors (CDKIs)) of cell cycle in purified Lin-BM cells. Earlier studies established that Cyclins and CDKIs are critical for the maintenance of proliferation vs. quiescence states in HSPCs^1,43^. Expression of CDKIs indicated normal levels of p27, but increased levels of p21 and p57 (**Figure 6A**) and of cyclins indicated normal levels of - *Ccnd2*, but a remarkable increase of *Ccna2*, (**Figure 6B**) in the Lin-cells of KO BM.

**Figure 6.**
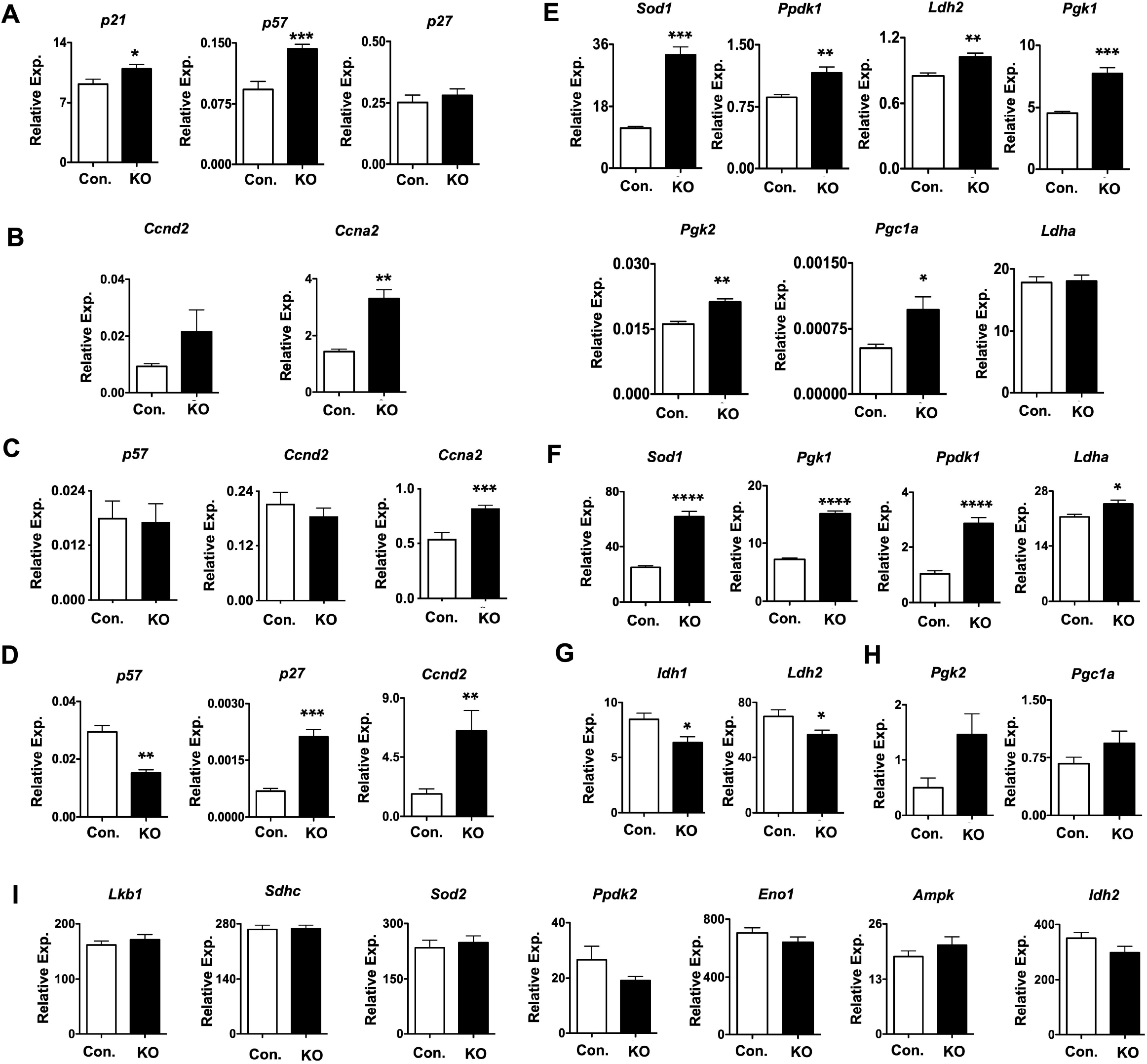
Deficiency of Trim28 affects cell cycle and metabolic genes in HSPCs. **A, B.** Real time PCR analysis of *p21*, *p57* and *p27* (**A**), and *Ccnd2* and *Ccna2* (**B**) mRNA levels in purified Lin^-^ cells of BM from KO (n=7) and control mice(n=8). Data are pool of 2 independent experiments. **C, D.** Real time PCR analysis of *p57*, *Ccnd2* and *Ccna2* mRNA levels in purified LSK subsets (**C**) and *p57*, *p27* and *Ccnd2* mRNA levels in purified LK subset of BM from KO (n=6) and control mice (n=6). Data are pool of 2 independent experiments. **E.** Real time PCR analysis of *Sod1*, *Ppdk1*, *Ldh2*, *Pgk1*, *Pgk2*, *Pgc1a and Ldha* mRNA levels in purified Lin^-^ cells of BM from KO (n=7) and control mice (n=8). Data are representative of 2 independent experiments. **F-I.** Real time PCR analysis of *Sod1*, *Pgk1*, *Ppdk1* and *Ldha* (**F**), *Idh1* and *Idh2* (**G**), *Pgk2* and *Pgc1a* (**H**), and *Lkb1*, *Sdhc*, *Sod2*, *Ppdk2*, *Eno1*, *Ampk* and *Idh2* (**I**) mRNA levels in purified LK subset of BM from KO (n=6) and control mice (n=6). Data are pool of 2 independent experiments. Expression levels of target genes were normalized to *Hprt/Gapdh/Actin* levels. All data represent mean ± SEM. Two-tailed Student’s *t* tests were used to assess statistical significance (*, P < 0.05; **, P < 0.01; ***, P < 0.001; ****, P < 0.0001).

To strengthen these findings, we purified LSK cells through flow cytometry and performed gene expression studies. Data indicated that the expression levels of *Ccna2*, but not *p57* and *Ccnd2*, were elevated in the LSK cells of KO BM (**Figure 6C**). On the other hand, analysis of purified LK cells revealed reduced levels of p57 and increased levels of *Ccnd2* and *p27* in the LK cells of KO BM (**Figure 6D**). To test if hyperproliferative phenotype of Trim28 deficient HSPCs is associated with altered metabolism, we analyzed the expression levels of genes involved in metabolic functions. Our data identified that expression levels of *Sod1*, *Ppdk1*, *Ldh2*, *Pgk1*, *Pgk2*, *Pgc1a* were elevated, and *Ldha* were normal in the Lin^-^ cells of KO mice (**Figure 6E**). Next, we studied purified LK cells and data revealed an upregulation of *Sod1*, *Ppdk1*, *Pgk1* and *Ldha* (**Figure 6F**), a down regulation of *Idh1* and *Ldh2* (**Figure 6G**), modestly increased expression levels, with no significance, of *Pgk2* and *Pgc1a* (**Figure 6H**), and normal expression levels of *Lkb1*, *Sdhc*, *Sod2*, *Ppdk2*, *Eno1*, *Ampk* and *Idh2* (**Figure 6I**) in the LK cells of KO BM. Overall, these data suggest that a loss of Trim28 results in hyperproliferation of HSPCs, possibly due to deregulated expression of key regulators of cell cycle and metabolism.

### Trim28 regulates transcription factors networks in HSPCs

Self-renewal, quiescence and differentiation of HSCs are governed mainly by the spatiotemporal expression of transcription factors^44,45^. Recent studies established that Trim28 has dual functions, as both an activator and a repressor, in regulating transcription^13^. To study if Trim28 deficiency leads to altered expression levels of transcription factors that promote HSC self-renewal and/or myeloid differentiation, we performed gene expression studies. Realtime PCR analysis on purified Lin^-^ BM cells suggested upregulated levels of *Gata1*, *Gata2*, *Gfi1* and *Meis1* (**Figure 7A**), and normal levels of *Klf1*, *Cebpa*, *Gfi1b*, *Pu.1 and Hoxa7* (**Figure 7B**) in KO BM. To assess the expression levels of transcription factors in myeloid progenitors, LK cells of BM were sorted, and data indicated increased expression of *Gata1*, *Gata2* and *Gfi1b* (**Figure 7C**), decreased expression of *Fli1*, *Meis1* and *Hoxa7* (**Figure 7D**), and normal expression levels of *Klf1*, *Gfi1* and *Pu.1* (**Figure 7E**) in the LK cells of KO BM. Next, we studied the expression levels of transcription factors in purified LSK cells. Realtime PCR studies indicated increased levels of *Gata1* and *Klf1* (**Figure 7F**) and normal levels of *Gfi1*, *Gfi1b*, *Gata2*, *Pu.1* and *Cebpa* (**Figure 7G**) in the LSK cells of KO BM.

**Figure 7.**
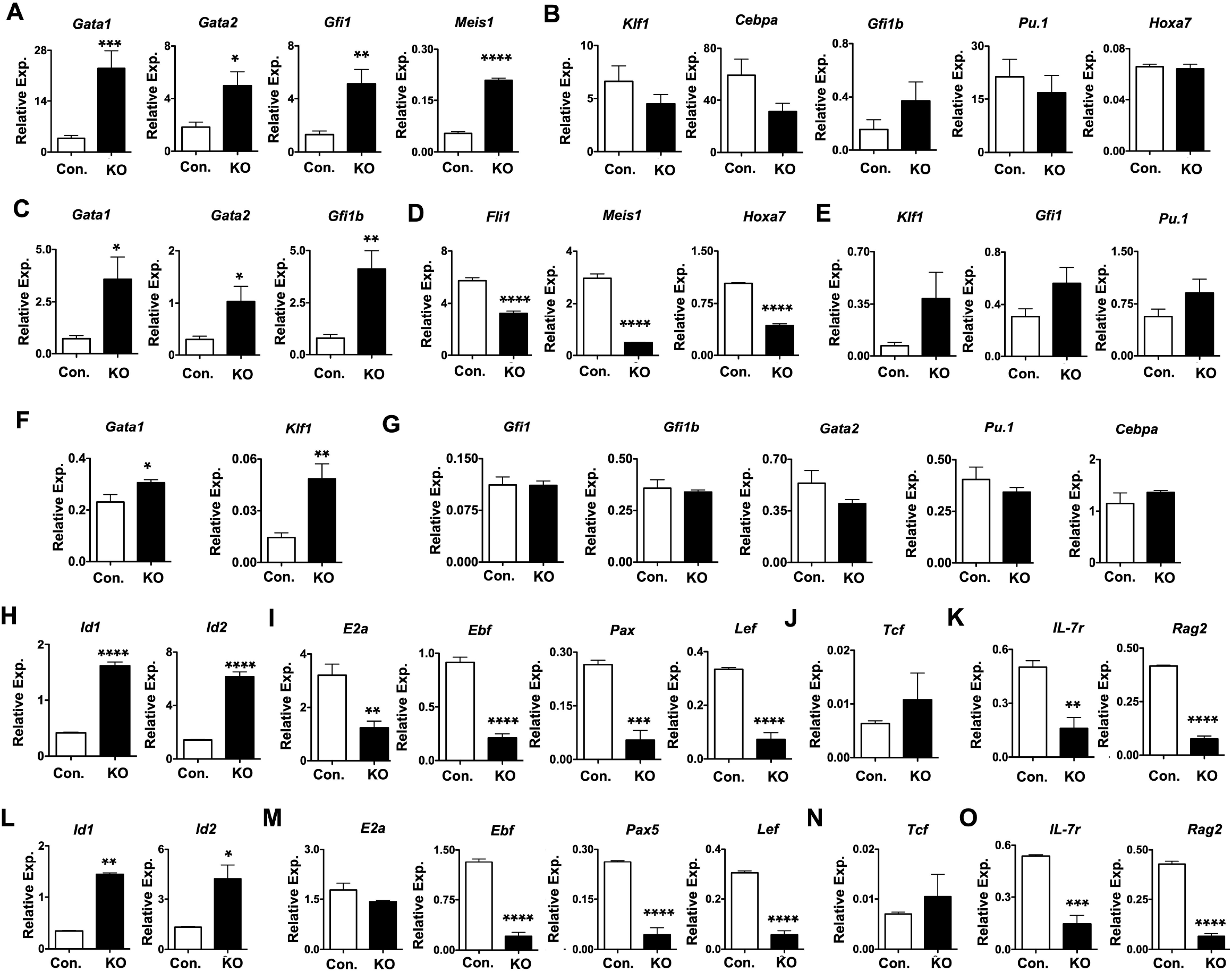
Deletion of Trim28 impairs transcription factors signatures in HSPCs. **A, B.** Real time PCR analysis of *Gata1*, *Gata2*, *Gfi1* & *Meis1* (**A**) and *Klf1*, *Cebpa*, *Gfi1b*, *Pu.1* & *Hoxa7* (**B**) mRNA levels in purified Lin-subset of BM from KO (n=7) and control mice (n=8). Data are pool of 3 independent experiments excluded data of *Meis1* that is representative of 2 experiments. **C-E.** Real time PCR analysis of *Gata1*, *Gata2* & *Gfi1b* (**C**), *Fli1*, *Meis1* & *Hoxa7* (**D**) and *Klf1*, *Gfi1* & *Pu.1* (**E**) mRNA levels in purified LK subset of BM from KO (n=6) and control mice (n=6). Data are pool of 2 independent experiments. **F, G.** Real time PCR analysis of *Gata1* & *Klf1* (**F**) and *Gfi1*, *Gfi1b*, *Gata2*, *Pu.1* & *Cebpa* (**G**) mRNA levels in purified LK subset of BM from KO (n=6) and control mice (n=6). Data are pool of 2 independent experiments. **H-K.** Real time PCR analysis of *Id1* & *Id2* (**H**), *E2a*, *Ebf*, *Pax5* & *Lef* (**I**), *Tcf* (**J**) and *Il7r* & *Rag2* (**K**) mRNA levels in purified Lin-subset of BM from KO (n=7) and control mice (n=8). Data are representative of 2 independent experiments. **L-O.** Real time PCR analysis of *Id1* & *Id2* (**L**), *E2a*, *Ebf*, *Pax5* & *Lef* (**M**), *Tcf* (**N**) and *Il7r* & *Rag2* (**O**) mRNA levels in purified CD19^+^B220^+^ subset of BM from KO (n=4) and control mice (n=5). Data are representative of 2 independent experiments. Expression levels of target genes were normalized to *Hprt/Gapdh/Actin* levels. All data represent mean ± SEM. Two-tailed Student’s *t* tests were used to assess statistical significance (*, P < 0.05; **, P < 0.01; ***, P < 0.001; ****, P < 0.0001).

Finally, we studied if the expression levels of transcription factors that dictate HSC differentiation into the lymphoid lineage are deregulated in the absence of Trim28. B cell differentiation in the BM is governed and strictly dependent on key transcription factors, such as E2a, Ebf and Pax5^46^. Realtime PCR studies on purified Lin-cells from KO BM indicated an upregulation of *Id1* and *Id2* (**Figure 7H**), a downregulation of *E2a*, *Ebf*, *Pax5* and *Lef1* (**Figure 7I**), and normal levels of *Tcf* (**Figure 7J**). Consistently, expression levels of key genes involved in lymphoid differentiation-*Il7r* and *Rag2*, were reduced in the Lin-cells of KO BM (**Figure 7K**). To corroborate and recapitulate these findings directly in B cells, we flow sorted CD19^+^B220^+^ B cells from the BM and performed Realtime PCR studies. Our data demonstrated that the expression levels of *Id1* and *Id2* were elevated (**Figure 7L**), of *E2a*, *Ebf*, *Pax5* and *Lef1* were downregulated (**Figure 7M**), and of *Tcf* were normal (**Figure 7N**) in the B cells of KO mice. In keeping with these findings, expression levels of *Il7r* and *Rag2* were strikingly reduced (**Figure 7O**) in purified B cells of KO mice. In essence, these data provide evidence that Trim28 protects the transcriptional landscape and transcription factors signatures of HSPCs and lymphoid lineage cells.

## Discussion

In the present study, we investigated the physiological roles of Trim28 in the maintenance and functions of HSCs. Our data reveal that a loss of Trim28 results in premature death due to pancytopenia, altered HSPC pool and aberrant HSPC functions. Our mechanistic studies established that Trim28 deficiency causes hyperproliferation of HSPCs, most likely due to deregulated expression of key regulators of cell cycle and metabolism. Furthermore, our data provided evidence that Trim28 is essential for maintaining the overall transcriptional landscape of HSPCs. In particular, Trim28 appears to regulate the expression of pivotal transcription factors that have been shown to be critical for the self-renewal, quiescence, multi-lineage differentiation and functions of HSCs^44,45^.

Trim28 has been implicated in a variety of physiologic and pathophysiologic processes including immune regulation, viral pathogenesis, cancer, tumor microenvironment and epithelial-to-mesenchymal transition^12,13^. Within the immune system, functions of Trim28 has been established in adaptive immune cells, both in B-^30^ and T-^31–33^ lineage cells. Even though Trim28 has been well recognized for its critical role in erythrocyte differentiation and functions^34,35^, their cellular and molecular functions in the self-renewal, quiescence and myeloid vs. lymphoid differentiation of HSCs remained elusive. While Miyagi et al. attempted to study the functions of Trim28 in HSPCs, most of their observations were based on conditional deletion of Trim28 using Tie2-Cre deleter strain^47^. Given the fact that Tie2-Cre mediated deletion of transgenes occurs mainly in endothelial cells^48,49^, and not specific to HSCs, and that endothelial cells play a crucial role in the maintenance of HSCs^4^, it is unclear as to what extent the observations of Miyagi et al. can be attributed to HSCs. More importantly, studies of Miyagi et al. were mainly based on fetal liver cells, which represents embryonic hematopoiesis^47^. In contrast, studies presented here is based on HSC specific deletion of Trim28 and on adult hematopoiesis. Due to these reasons, unsurprisingly, our observations and conclusion are different from that of Miyagi et al., even though there are some overlapping findings between our studies.

Emerging studies have unequivocally established that Trim28 is a highly versatile multidomain protein^12^ with pleiotropic functions. Despite being recognized as a powerful suppressor of transcription^50^, Trim28 has been shown to function as a transcriptional activator^13^. Even though Trim28 has been shown to bind promoter/regulatory regions of over 3,000 genes^27^, protein-protein interactions have been identified to play a central role in the transcriptional regulation function of Trim28^51^. Furthermore, Trim28 exhibits transcriptional co-repressor activities by forming stable repressive complexes with key proteins such as, KRAB-domain zinc finger DNA binding factors, SETDB1, HP1, NuRD deacetylase complex, and histone H3K9 methyltransferase ESET^12,50^. While the pivotal functions of Trim28 in the transcriptional regulation of target genes have been well established, precise mechanisms through which Trim28 regulates its target genes remain unclear^12^. Data from the present study established that Trim28 deficiency causes major alterations in the gene expression profiles, including key transcription factors and cell cycle regulators, in HSPCs. However, currently, it is unclear if these molecular changes are directly due to defective transcriptional repression and/or activation by Trim28. Furthermore, it needs to be determined as how many of these identified transcription factors are direct transcriptional targets of Trim28 in HSPCs. Of note, previous studies identified abnormal microRNA expression profiles in Trim28 deficient erythroid lineage cells^34^. Given the critical functions of microRNAs^52^ and its machinery^53^ in hematopoiesis, we cannot rule out the possible involvement of microRNAs in the phenotype of Trim28 deficient HSPCs.

Post-translational modifications of proteins, especially ubiquitylation, have been shown to play pivotal roles in hematopoiesis, particularly in the maintenance of HSCs^3^. A series of studies from our lab demonstrated that a deficiency of c-Cbl (a member of RING family of E3 ubiquitin ligases) results in loss of HSC pool and functions^54^, age related myeloproliferation and lymphopenia^55^, and the onset of acute myeloid leukemia^56^. Consistently, our studies demonstrated that loss of Itch (a member of HECT domain family of E3 ligases) caused abnormal hematopoiesis and defective HSC functions^57^. Recent studies from our lab established that a deficiency of A20 (a ubiquitin editing enzyme) causes severe HSC defects and functions, which ultimately leads to pancytopenia and premature death^37,58,59^. Interestingly, Trim28 protein possesses a RING finger domain and has been identified to exert E3 ubiquitin ligase functions on substrates including p53, AMPK and CDK9 proteins^13,60^. However, to date, the E3 ligase functions of Trim28 in HSCs remain largely unidentified. Of note, the hematopoietic phenotype, such as pancytopenia, HSC pool and functional defects, and premature death, of Trim28 deficient mice greatly resembles the phenotype of our previously described mouse models with ubiquitylation defects^37,58,59^. Based on these, it is tempting to speculate that the E3 ubiquitin ligase functions of Trim28 is essential for the maintenance and functions of HSPCs. Even though we do not provide any evidence in favor of this hypothesis in this manuscript, an in-depth mechanistic study to identify the E3 ligase functions of Trim28 is an active area of our investigation.

In essence, our studies provide several novel insights, especially on the roles of Trim28 in the maintenance of HSPC pool and their differentiation into myeloid and lymphoid lineages. More importantly, our studies identify candidate hematopoietic transcription factors that are deregulated in the absence of Trim28 in HSPCs.

## Materials and Methods

### Mice

Trim28 Floxed mice^20^ and Vav-iCre (B6.Cg-Commd10^Tg(Vav1-icre)A2Kio^/J) mice were purchased from the Jackson Laboratory. CD45.1 congenic animals were purchased from the National Cancer Institute. The Institutional Animal Care and Use Committee approved all mouse experiments.

#### Cell preparation

Mice were analyzed between 3 and 4 weeks after birth. RBCs were lysed with ammonium chloride (STEMCELL Technologies). Trypan blue (Amresco)–negative cells were counted as live cells.

#### Flow cytometry and Cell Sorting

Cells were analyzed by flow cytometry with Attune Nxt (Thermofisher) and FlowJo software (Tree Star). Cells were sorted using FACS Aria and Magnetic associated cell sorting (Miltenyi). The following monoclonal antibodies were used: anti-CD34 (RAM34), anti-CD45.1 (A20), anti-CD45.2 (104), anti-CD48 (HM48-1), anti-CD117 (2B8), anti-Flt3 (A2F10.1), anti-Sca-1 (D7), anti-B220 (RA3-6B2), anti-CD19 (1D3), anti-CD3 (145-2C11), anti-CD4 (GK1.5), anti-CD8 (53-6.7), anti-CD11b (M1/70), anti– Gr-1 (RB6-8C5) and anti-Ter119 (TER119; from BD Biosciences); anti-CD150 (TC15-12F12.2) from Biolegend; anti-CD16/32 (93) from eBioscience. Cells incubated with biotinylated monoclonal antibodies were incubated with fluorochrome-conjugated streptavidin–peridinin chlorophyll protein–cyanine 5.5 (551419; BD), streptavidin-allophycocyanin-Cy7 (554063; BD), streptavidin-super bright 650 (Biolegend). In all the FACS plots, indicated are the percentages (%) of the gated fraction. The gating strategies for multi-lineage and HSPC analyses are indicated in **Supplemental Figures 5 & 6**.

#### Bone marrow and serial transplantation experiments

1 × 10^6^ of bone marrow cells were injected into lethally irradiated (10 Gy) congenic (CD45.1^+^) recipient mice. For competitive-repopulation experiments, 5 × 10^5^ BM cells from either control or KO mice were mixed with 5 × 10^5^ of WT (CD45.1^+^) BM cells (to obtain a ratio of 1:1) and were injected into lethally irradiated congenic WT (CD45.1^+^) recipient mice. For serial transplantation assays, 1 × 10^6^ of bone marrow cells were injected into lethally irradiated (10 Gy) WT congenic (CD45.1^+^) recipient mice. After 12 weeks of transplantation, 1 × 10^6^ BM cells of primary recipients were injected into lethally irradiated WT congenic secondary recipients.

### BrdU incorporation and Ki67 studies

For *in-vivo* bromodeoxyuridine (BrdU) assay, 1 mg BrdU (BD) was injected intraperitoneally. After 24 hours of injection, mice were sacrificed and bone marrow cells were stained for BrdU, following the BrdU Flow Kit manufacturer’s instructions (BD Pharmingen). For Ki67 analysis, BM cells were stained for cell surface markers, fixed, and permeabilized with BD Fix/Perm kit. Cells were stained with anti-Ki67—APC (BD558615) for 30 min on ice and analyzed by flow cytometry.

#### RNA extraction, PCR, and real-time PCR

Total RNA was isolated using RNeasy Mini kit or RNeasy Micro kit (QIAGEN). cDNA was synthesized with Oligo(dT) primer and Superscript IVReverse Transcriptase (Thermo Fisher Scientific). PCR was performed with T100 thermal cycler (Bio-Rad Laboratories) and TSG Taq (Lamda Biotech). Real-time PCR was performed in duplicates with a CFX-connect real-time PCR system (Bio-Rad Laboratories) and SsoAd-vanced SYBR Green Supermix (Bio-Rad Laboratories) according to the manufacturer’s instructions. Relative expression was normalized to the expression levels of the internal control (housekeeping gene) HPRT/GAPDH.

#### Western blot analysis

For immunoblot analyses, cells were lysed with cell lysis buffer (Cell Signaling Technology) with protease inhibitor cocktail (Complete; Roche) and 1 mM PMSF (Santa Cruz Biotechnology, Inc.). Cell lysates were boiled with sample buffer (NuPAGE; Life Technologies) containing 1% 2-Mercaptoethanol (Sigma-Aldrich). Proteins were subjected to 8–12% SDS-PAGE and transferred to PVDF membranes (Bio-Rad Laboratories). The membranes were blocked with 5% skim milk and then treated with Trim28 primary and anti–rabbit IgG (Cell Signaling Technology) secondary antibodies, respectively.

#### Statistics

Data represent mean and s.e.m. Two-tailed student’s t and non-parametric test (Mann Whitney) tests were used to assess statistical significance (*P < 0.05, **P<0.01, *** < 0.001). For survival curve analysis, log rank test was used to assess statistical significance (****P< 0.0001).

## Author Contributions

GS performed all experiments, collected the data, analyzed the data, and prepared figures. CR designed research, analyzed, and interpreted the data, prepared figures, and wrote and corrected the manuscript. Both authors contributed to the article and approved the submitted version.

