## Supplemental Figures for "Trim28 plays an indispensable role in maintaining functions and transcriptional integrity of hematopoietic stem cells"

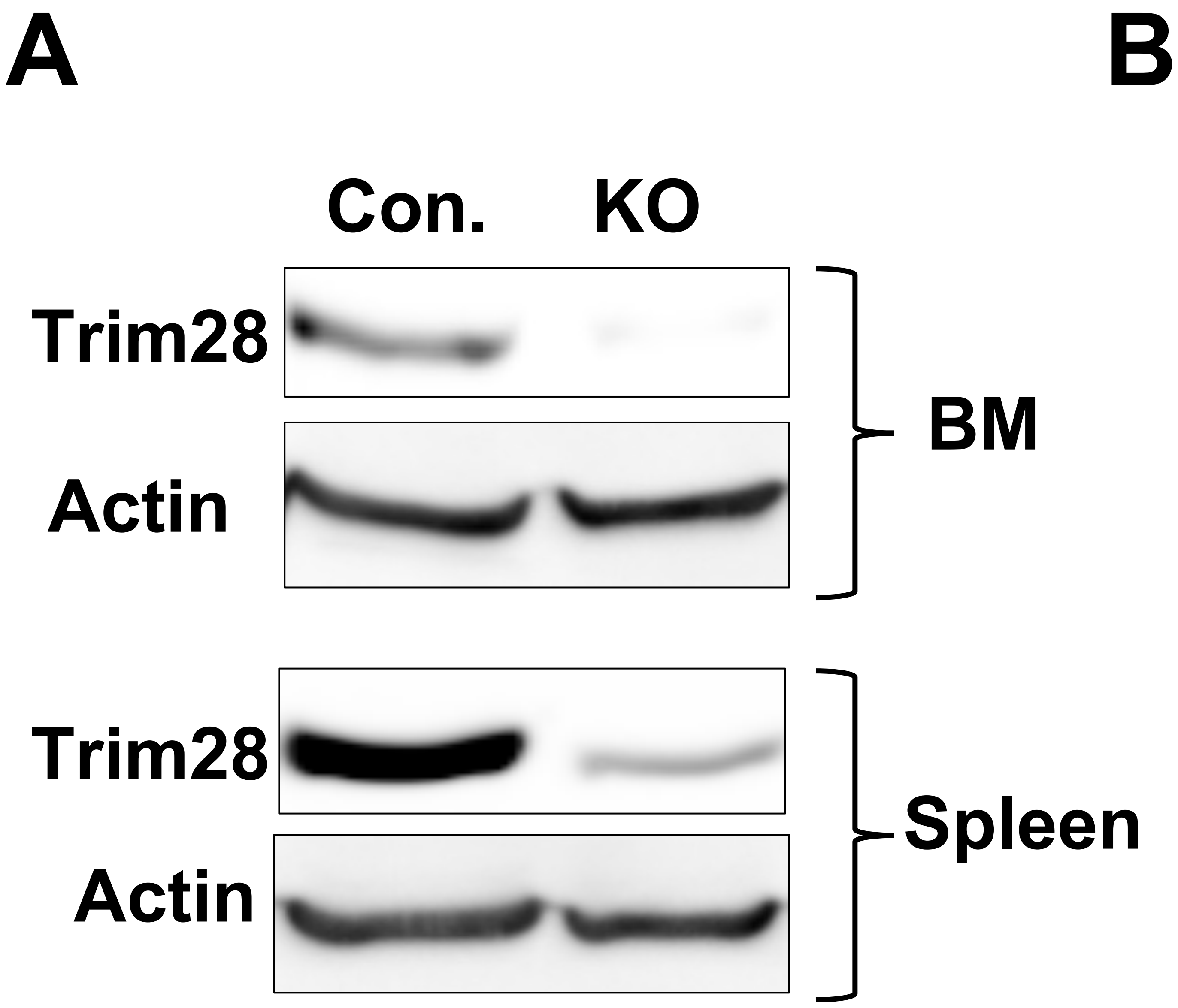

**B**

| Litters | No of mice | Trim28 F/+ | Trim28 F/F | Trim28 F/+ Vav cre | Trim28 F/F Vav cre |
|---|---|---|---|---|---|
| Litter 1 | 19 | 6 | 3 | 10 | 0 |
| Litter 2 | 26 | 7 | 8 | 10 | 1 |
| Litter 3 | 48 | 13 | 14 | 14 | 7 |
| Litter 4 | 15 | 6 | 1 | 7 | 1 |
| Litter 5 | 21 | 5 | 7 | 6 | 3 |
| Litter 6 | 24 | 5 | 4 | 15 | 0 |
| Litter 7 | 22 | 9 | 4 | 9 | 0 |
| Litter 8 | 28 | 9 | 8 | 11 | 0 |
| Litter 9 | 26 | 9 | 13 | 3 | 1 |
| Litter 10 | 11 | 4 | 6 | 1 | 0 |
| Total | 240 | 73 | 68 | 86 | 13 |

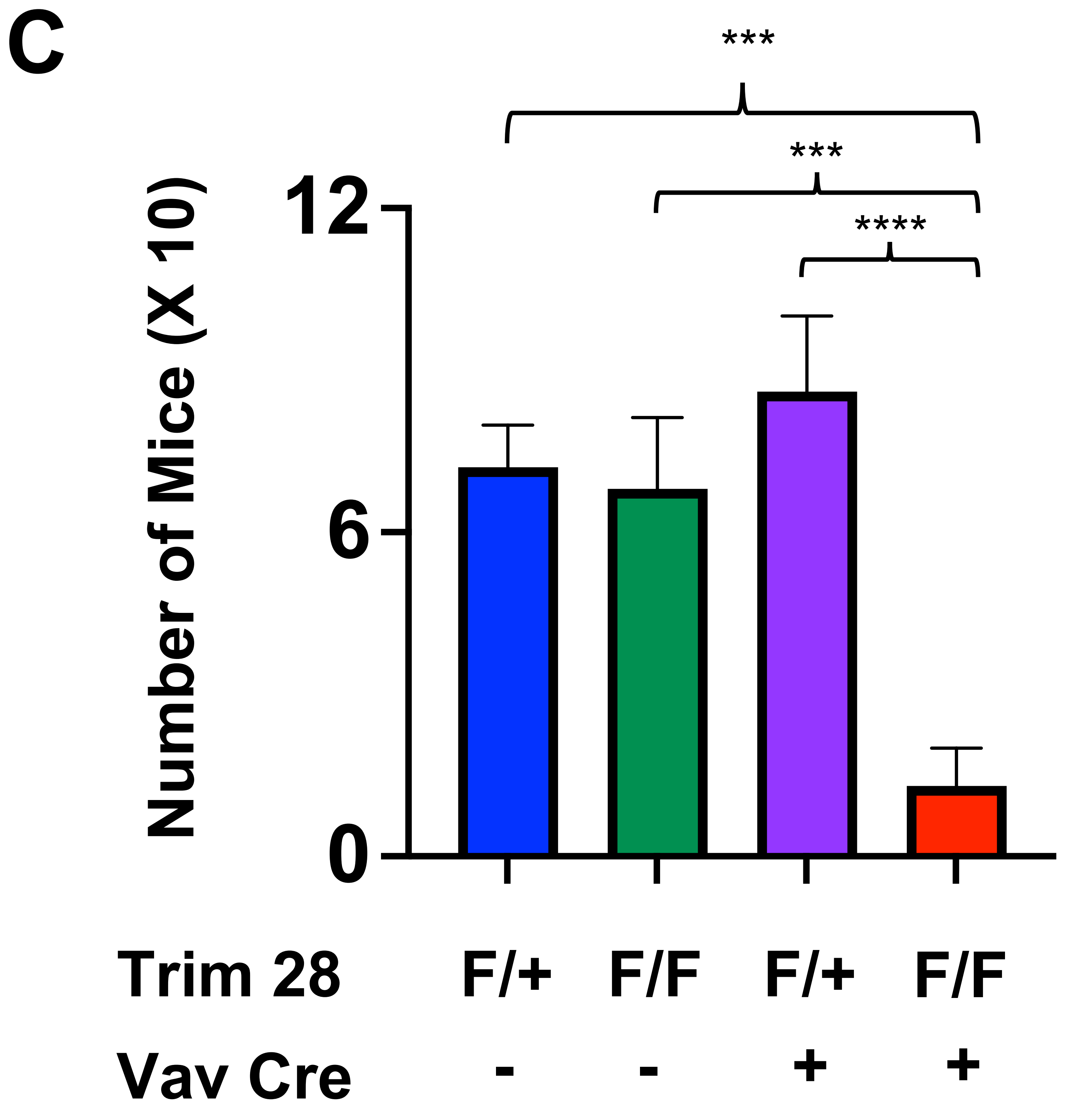

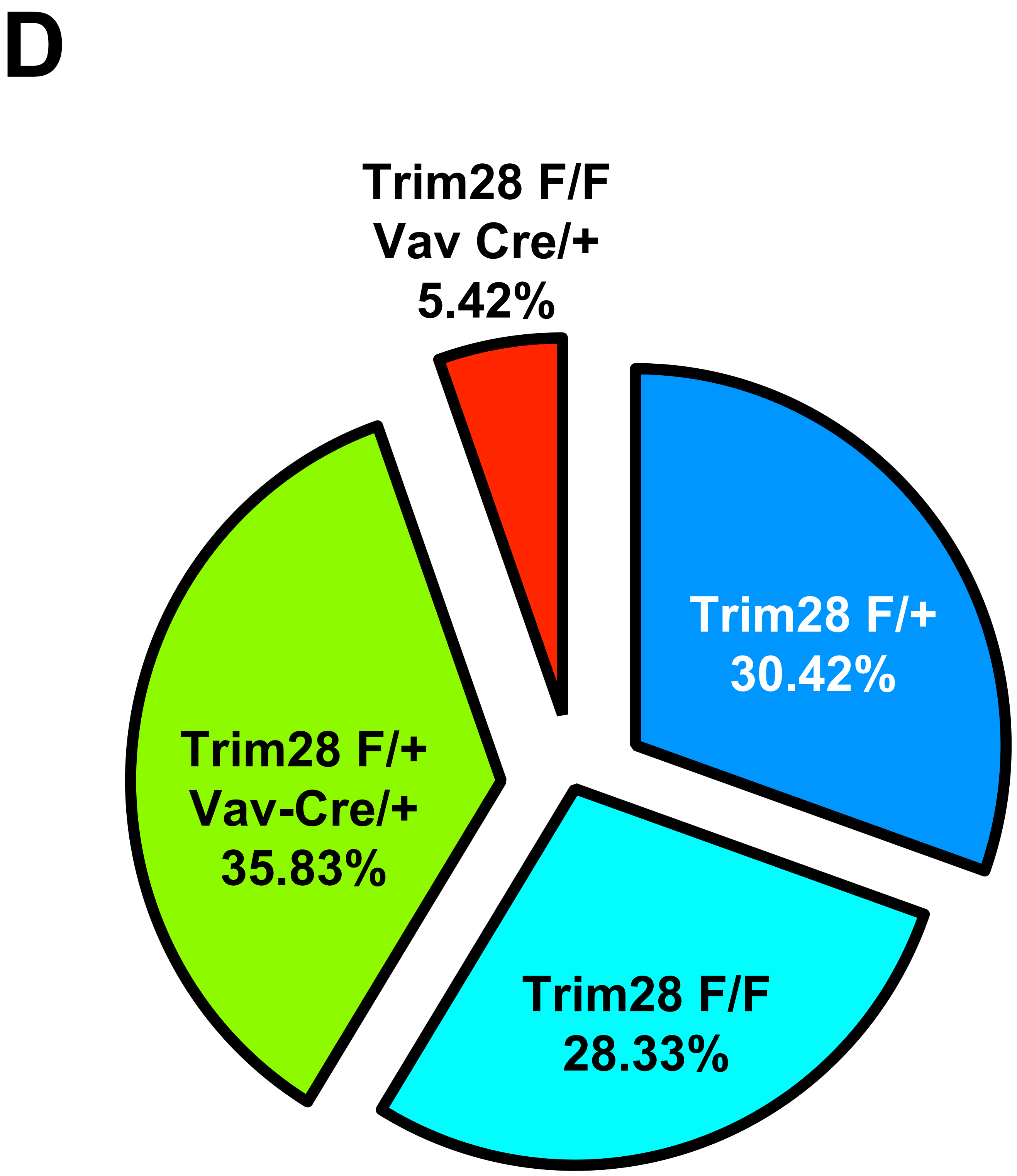

Supplemental Figure 1.

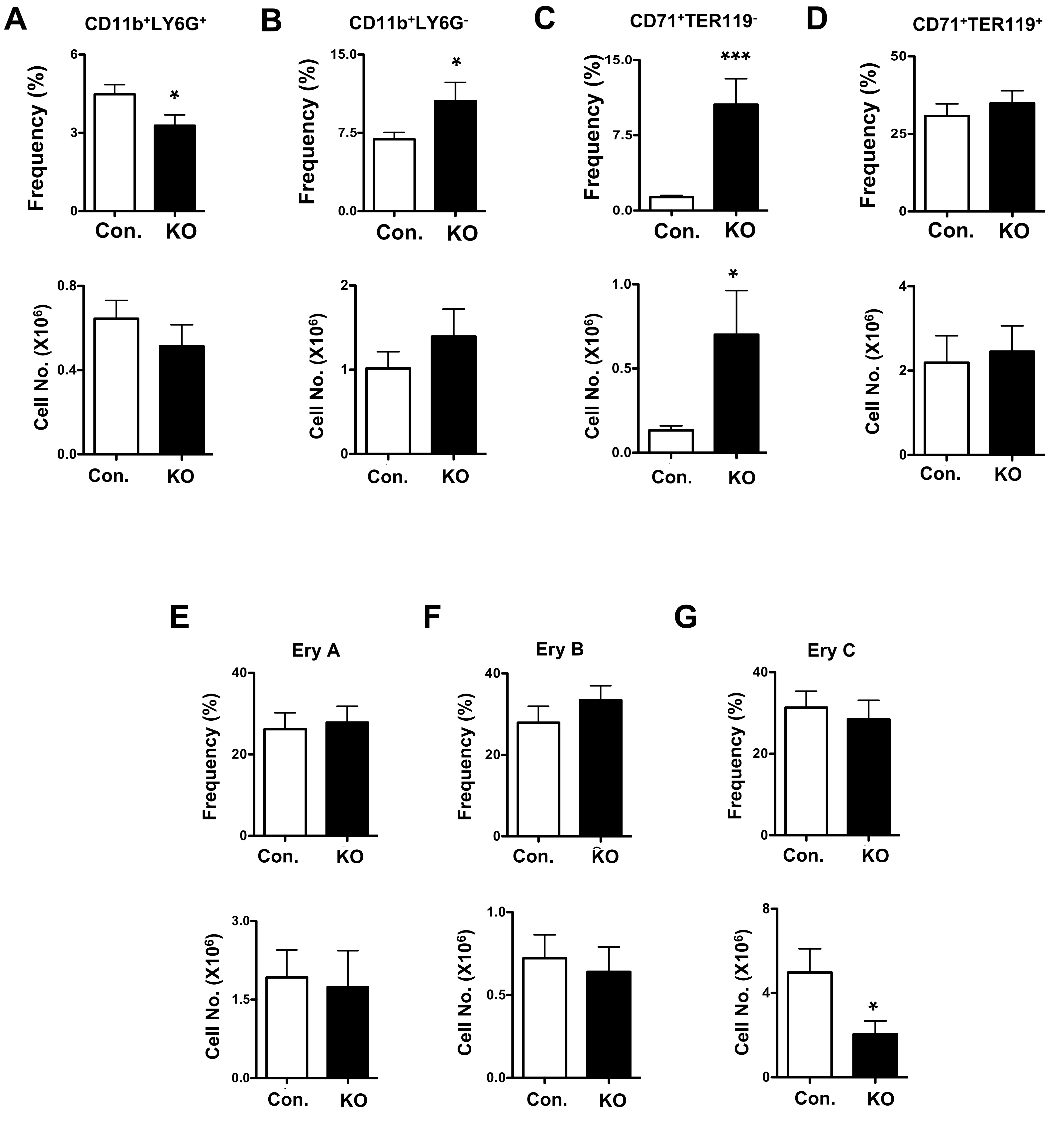

### Supplemental Figure 2.

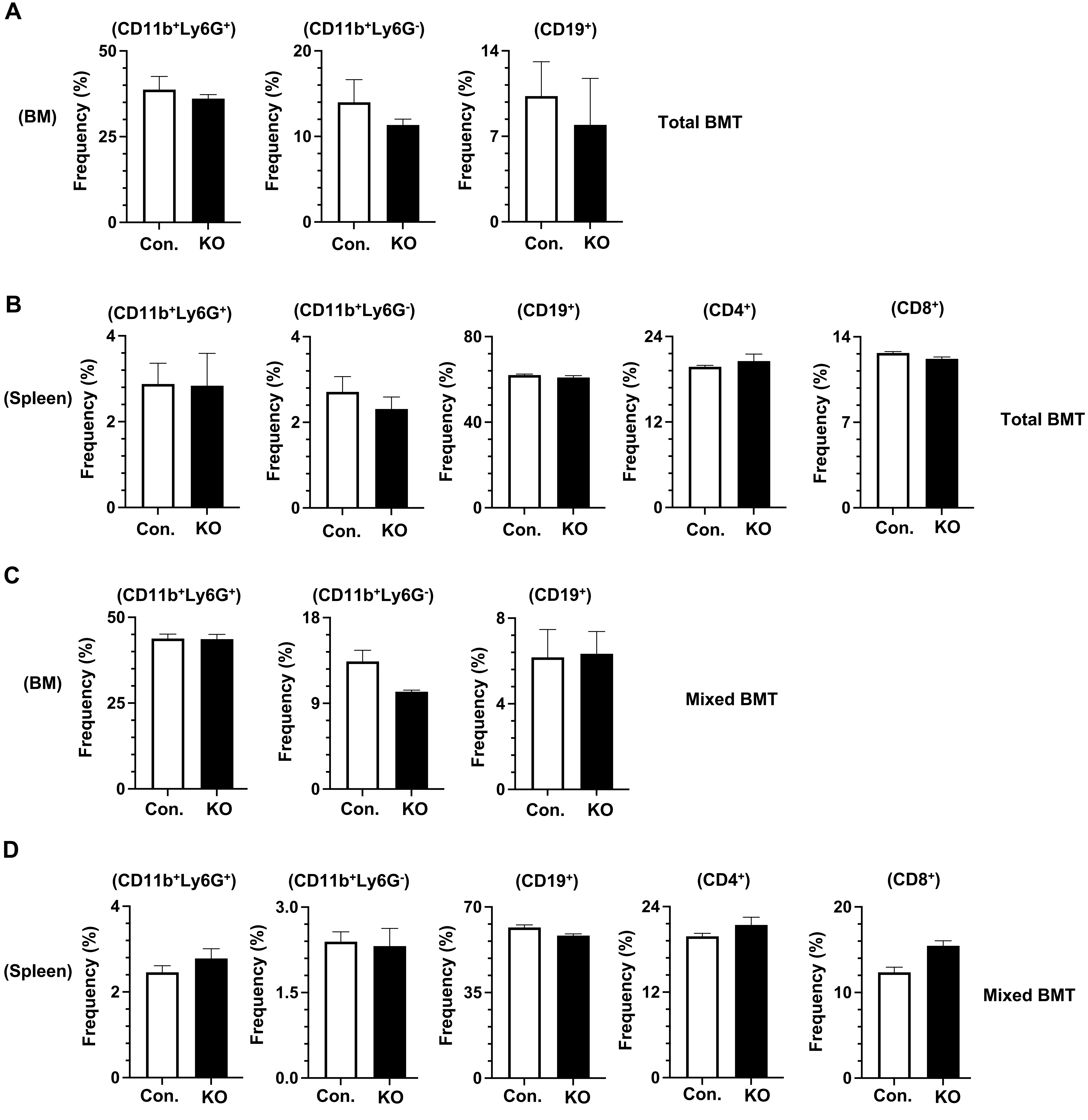

**Supplemental Figure 3.**

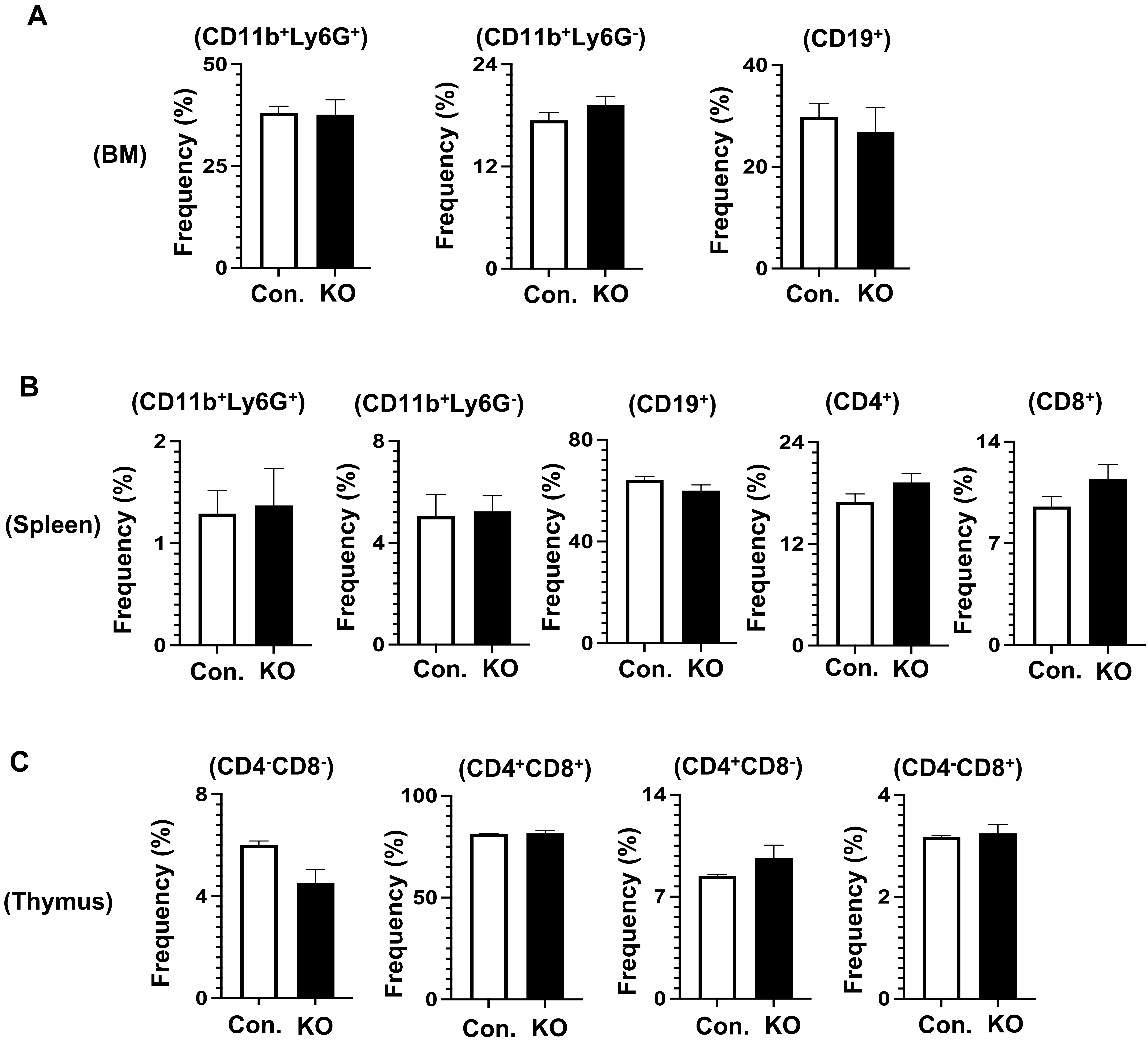

Supplemental Figure 4.

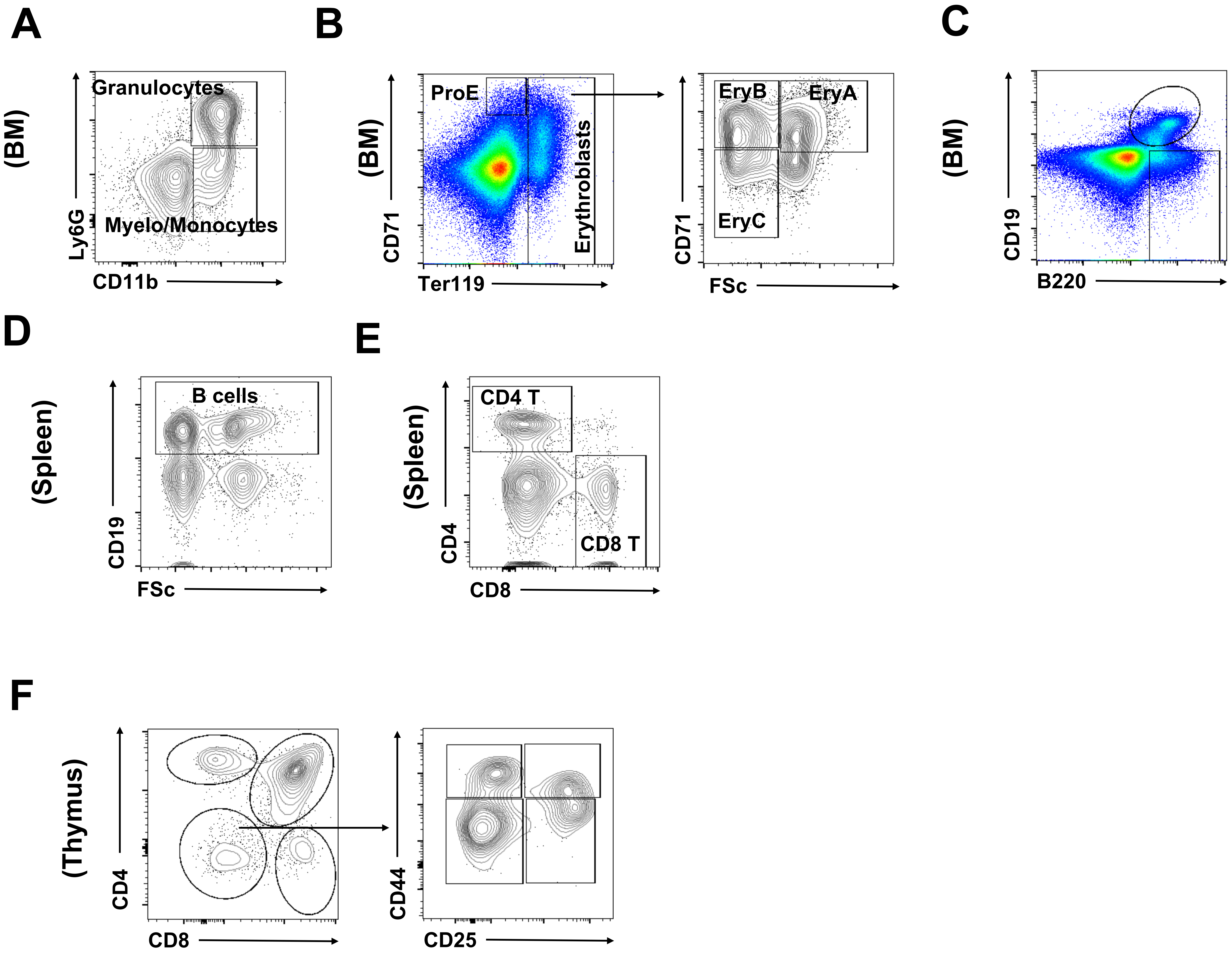

**Supplemental Figure 5.**

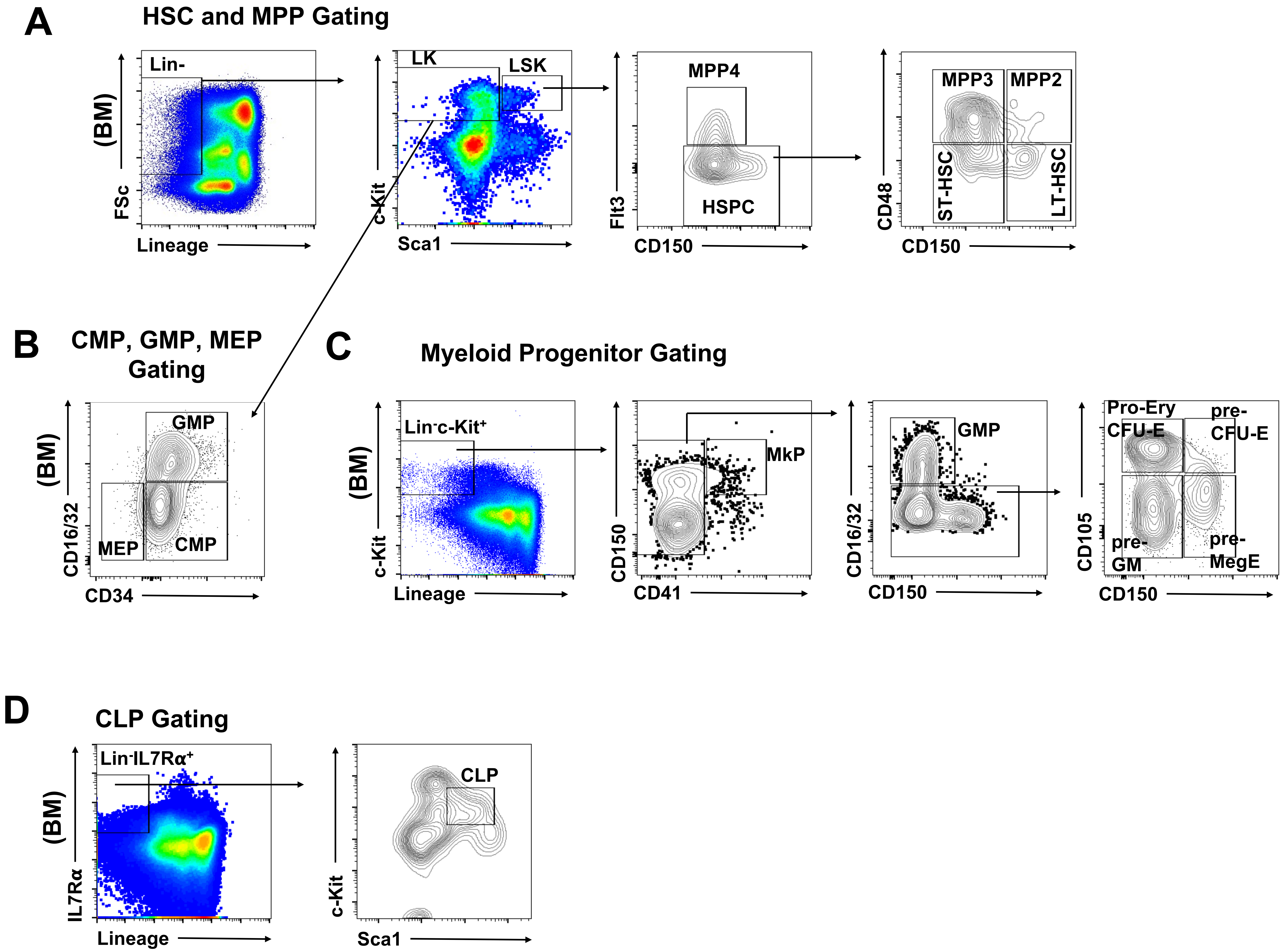

**Supplemental Figure 6.**
