## Supplemental Figure Legends for "Trim28 plays an indispensable role in maintaining functions and transcriptional integrity of hematopoietic stem cells"

**Supplemental Figure 1.**

**A.** Western blot analysis of Trim28 expression in the BM and spleen of KO and Con mice.

**B-D.** Details of genotyping of litters obtained from Trim28 breeding pairs. A total of 240 animals from 10 independent litters were taken for the study.

**Supplemental Figure 2.**

**A-D.** Frequencies (**top**) and absolute numbers (**bottom**) of granulocytes (**A**), myelo/monocytes (**B**), proerythroblasts (**C**) and total erythroblasts (**D**) in the spleen of KO (n=21) and Con. (n=26) mice between 3 and 4 weeks of birth.

**E-F.** Frequencies (**top**) and absolute numbers (**bottom**) of Immature erythroblasts (EryA) (**E**), less-mature erythroblasts (EryB) (**F**), and most-mature erythroblasts (EryC) (**G**) in the spleen of KO (n=21) and Con. (n=26) mice between 3 and 4 weeks of birth.

All data represent mean ± SEM. Two-tailed Student's *t* tests were used to assess statistical significance (*, P < 0.05; **, P < 0.01; ***, P < 0.001; ****, P < 0.0001).

**Supplemental Figure 3.**

**A.** Frequencies of total granulocytes (**left**), myelo/monocytes (**middle**) and B cells (**right**) in the BM of WT recipients (n= 9), after 24 weeks of total-BMT.

**B.** Frequencies of total granulocytes (**far left**), myelo/monocytes (**left**), B cells (**center**), $CD4^{+}$T cells (**right**) and $CD8^{+}$T cells (**far right**) in the spleen of WT recipients (n= 9), after 24 weeks of total-BMT.

**C.** Frequencies of total granulocytes (**left**), myelo/monocytes (**middle**) and B cells (**right**) in the BM of WT recipients (n=14), after 24 weeks of mixed-BMT.

**D.** Frequencies of total granulocytes (**far left**), myelo/monocytes (**left**), B cells (**center**), $CD4^{+}$T cells (**right**) and $CD8^{+}$T cells (**far right**) in the spleen of WT recipients (n= 14), after 24 weeks of mixed-BMT.

**Supplemental Figure 4.**

**A.** Frequencies of total granulocytes (**left**), myelo/monocytes (**middle**) and B cells (**right**) in the BM of WT recipients (n= 6), after 16 weeks of secondary-BMT.

**B.** Frequencies of total granulocytes (**far left**), myelo/monocytes (**left**), B cells (**center**), $CD4^+$T cells (**right**) and $CD8^+$T cells (**far right**) in the spleen of WT recipients (n= 6), after 16 weeks of secondary-BMT.

**C.** Frequencies of DN (**far left**), DP (**left**), SP4 (**right**) and SP8 (**far right**) subsets in the thymus of WT recipients (n= 6), after 16 weeks of secondary-BMT.

**Supplemental Figure 5. Gating scheme for multi-lineage analysis.**

**A.** Gating scheme for granulocytes and myelo/monocytes of the BM.

**B.** Gating scheme for proerythrocytes and erythroblast subsets of the BM.

**C.** Gating scheme for B cells of the BM.

**D.** Gating scheme for B cells of the spleen.

**E.** Gating scheme for $CD4^+$ and $CD8^+$ T cells of the spleen.

**F.** Gating scheme for thymic subset analysis.

**Supplemental Figure 6. Gating scheme for HSPC analysis.**

**A.** Gating scheme for $Lin^-$, LK, LSK, LT-HSC, ST-HSC, MPP2, MPP3 & MPP4 analysis of the BM.

**B.** Gating scheme for CMP, GMP & MEP analysis of the BM.

**C.** Gating scheme for pre-GMP, GMP, pre-MegE, MkP, pre-CFU-E & pro-Ery/ CFU-E analysis of the BM.

**D.** Gating scheme for CLP analysis of the BM.
